# Interleukin 6 drives durable T cell-mediated immunity to pancreatic cancer

**DOI:** 10.1101/2024.09.26.615308

**Authors:** Paige C. Arneson-Wissink, Alexandra Q. Bartlett, Heike Mendez, Xinxia Zhu, Peter R. Levasseur, Parham Diba, Namratha Turuvekere Vittala Murthy, Jessica Dickie, Matthew McWhorter, Antony Jozic, Yulia Eygeris, Gaurav Sahay, Katelyn T. Byrne, Gregory D. Scott, Robert Eil, Aaron J. Grossberg

**Author notes:** **Corresponding Author**: Aaron J. Grossberg, 2730 S Moody Ave, Oregon Health and Science University, Portland, OR, 97201.

## Abstract

**Background & Aims:** Tumor immune resistance is recognized as a contributor to low survivorship in pancreatic ductal adenocarcinoma (PDAC). The inflammatory cytokine interleukin-6 (IL-6) promotes polarization of CD4 T cell populations away from immune tolerance, and induces differentiation of cytotoxic CD8 T cells. This work aims to test whether IL-6 could stimulate an anti-tumor response in PDAC

**Methods:** We overexpressed IL-6 in multiple Kras^G12D/+^, Tp53^R172H/+,^ Pdx1-Cre (KPC) cell lines, which were orthotopically implanted in mice (OT-PDAC^IL6^). We followed mouse survival and measured tumor growth, tumor histology, and plasma IL-6 at 5 and 10 days after tumor implantation. We measured tumor immune cell infiltration via flow cytometry and histology. We used antibody-based T cell depletion and secondary tumor implantation rechallenge to test the dependency of the durable immune reaction on T cells. We use lipid nanoparticle (LNP)-based delivery of IL-6 mRNA to the pancreas as an orthogonal approach for testing the effect of elevated IL-6 in the tumor microenvironment on anti-tumor T cell invasion.

**Results:** Improved survival occurred in all instances of OT-PDAC^IL6^, with one cell line (KxPxCx) reproducibly resulting in long-term recurrence-free survival. With KxPxCx cells, circulating IL-6 was 100-fold higher in OT-KxPxCx^IL6^ than in OT-KxPxCx^parental^ mice. Flow cytometry revealed increased T cells and NK cells, and decreased T regulatory cells, and we observed significantly increased lymphoid aggregates in OT-KXPXCX^IL6^ as compared to OT--KxPxCx^parental^ tumors. Antibody-based CD4^+^ and CD8^+^ T cell depletion prevented tumor clearance and completely abolished the survival advantage in OT-KxPxCx^IL6^ mice. The anti-tumor immune response to OT-KxPxCx^IL6^ rendered mice immune to re-challenge with OT-KxPxCx^parental^ tumors. LNP delivery of IL-6 to the pancreas elevated systemic IL-6 levels ∼50 fold, lowered tumor burden, and increased anti-tumor T cell phenotypes.

**Conclusions:** Locally high IL-6 concentrations potently enhance the T cell-mediated anti-tumor response to PDAC.

**SYNOPSIS:** Interleukin-6 induces rapid and durable T cell-driven immune clearance of pancreatic ductal adenocarcinoma. The anti-tumor immune microenvironment is hallmarked by increased lymphoid aggregate formation, increased CD4 T cell abundance, and decreased Treg abundance.

## INTRODUCTION

Pancreatic ductal adenocarcinoma (PDAC) is associated with an immunosuppressive microenvironment, characterized by low numbers of infiltrating T cells and relatively higher numbers of immunosuppressive T regulatory cells (Tregs)^1, 2^. Despite clinical trials that have established the safety of immunotherapies in PDAC, the immune profile of PDAC limits the clinical efficacy of these interventions^3^. Conversely, increased T cell infiltration with low Treg infiltration is associated with improved patient survival ^5^. Strategies that promote T cell activation, such as agonizing CD40 antibodies, increase tertiary immune structure formation and benefit mouse survival in pre-clinical models^4^. A small but growing body of literature suggests that tumor-localized expression of immunomodulatory cytokines, such as IL-6, IL-12, CXCL13, and CCL21 can also improve immune reactivity to tumors by driving localization and differentiation of tumor-infiltrating lymphocytes, which accumulate in tertiary lymphoid structures or lymphoid aggregates^5–7^.

Interleukin-6 (IL-6) signaling is associated with reduced survival in patients with PDAC attributed to the promotion of tumorigenesis and metastasis ^8–11^. Elevated circulating IL-6 also associates with sarcopenia in PDAC patients, and we previously demonstrated that host intrinsic IL-6 signaling is necessary for mice to develop PDAC cachexia^12^. Published literature shows that loss of IL-6 attenuates PDAC development, metastasis, and cachexia onset ^12–14^. However, IL-6 is also well-established as an immunomodulatory cytokine and contributes to polarization of CD4 T cell populations, differentiation of cytotoxic CD8 T cells, inhibition of inducible Treg development, and expansion and anti-tumor activity in chimeric antigen receptor (CAR) T cell therapies^15–17^. Leveraging the immunomodulatory properties of IL-6 may promote anti-tumor immunity in PDAC.

In the current study, we sought to investigate the effect of IL-6 overexpression on immune clearance of PDAC and cachexia development. We used two independent modalities to elevate IL-6 expression in the pancreas: stable IL-6 overexpressing PDAC cells (PDAC^IL6^) from PDAC^parental^ (KPC) cell lines^18^, and lipid nanoparticles (LNPs) containing IL-6 mRNA. We found that PDAC^IL6^ cells produced extremely high levels of IL-6 and induced severe cachexia within days of orthotopic implantation (OT-PDAC^IL6^). We also observed dramatic changes in tumor growth dynamics, with one cell line eliciting complete tumor clearance and long-term, recurrence-free mouse survival. IL-6 LNPs targeted the pancreas, caused high levels of circulating IL-6, and limited PDAC tumor growth. We then pursued a series of studies to understand the immune response to high intra-tumoral IL-6. The work shown here presents IL-6 overexpression as a driver of T cell infiltration in PDAC, which poises mice for a favorable anti-tumor immune response.

## RESULTS

### Tumor-specific IL-6 overexpression induces spontaneous tumor clearance and cachexia recovery

We developed PDAC^IL6^ cells from the KPC PDAC cell line KxPxCx, using ecotropic retroviral transduction. We first observed that mice implanted with OT-KxPxCx^IL6^ tumors experienced nearly 100% survival without reaching terminal endpoints (**Figure 1A**). OT-KxPxCx^IL6^ mice euthanized approximately 10 days after all OT-KxPxCx^parental^ had significantly lower tumor/pancreas mass, and no detectable IL-6 in the plasma (**Figure 1B-C**). Deeper histological assessment of tumors showed that both OT-KxPxCx^IL6^ and OT-KxPxCx^parental^ developed poorly differentiated, infiltrative carcinoma by five days (**Figure 1D-F**). OT-KxPxCx^IL6^ tumors were slightly smaller in mass and histological area than OT-KxPxCx^parental^ tumors (**Figure 1F-G**). By 12 days, OT-KxPxCx^parental^ mice had reached humane euthanasia endpoint, with carcinoma covering approximately 80% of total tissue area, in contrast, OT-KxPxCx^IL6^ tissue was completely devoid of tumor (**Figure 1E,G, Table S1**). We confirmed this result molecularly using qPCR for the codon-optimized *Il6* transgene, which was not detected in OT-KxPxCx^IL6^ whole pancreas tissue after five days (**Figure 1H, Figure S1**). While lung and liver metastases were detected by *Il6* transgene qPCR at five days, there were no metastases detected at 12 days (**Figure S2**). Spontaneous tumor clearance also occurred when KxPxCx^IL6^ cells were implanted subcutaneously, indicating that the phenomenon is not specific to intrapancreatic administration (**Figure S3**).

**Figure 1:**
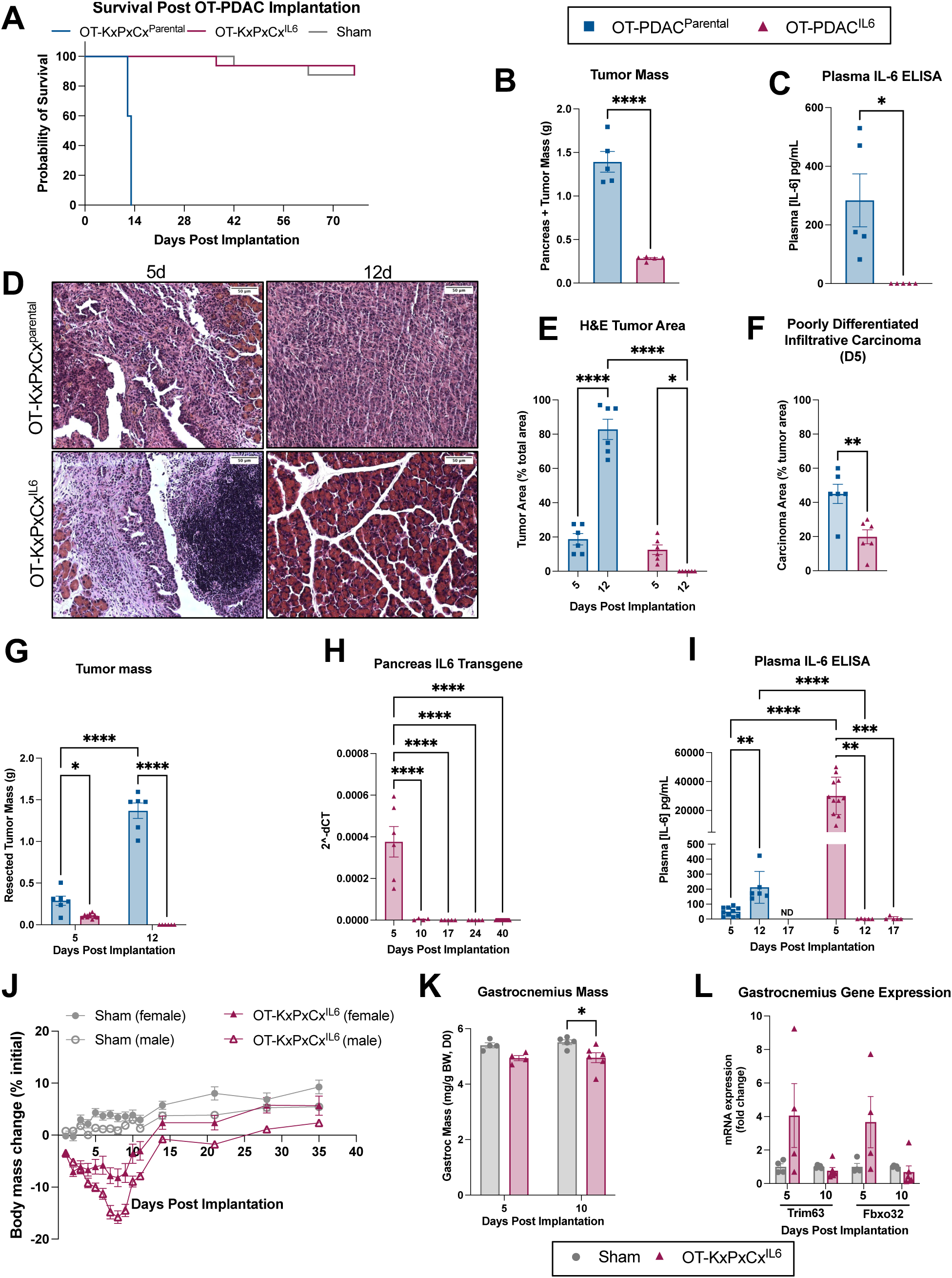
OT-KxPxCx^IL6^ results in complete tumor clearance and body mass recovery. (A) Survival comparison of OT-KxPxCx^Parental^ and OT-KxPxCx^IL6^. N = 8 female, 8 male Sham and OT-KxPxCx^IL6^, 1 female, 4 male OT-KxPxCx^parental^ mice. Statistically tested with Log-rank (Mantel-Cox) test. (B) Tumor mass at humane endpoint (OT-KxPxCx^Parental^). Pancreas mass of OT-KxPxCx^IL6^ euthanized at a pre-determined endpoint of 24 days (did not meet humane euthanasia criteria); tumor tissue was not present. Statistically tested with 2-way ANOVA main effects only with Tukey correction for multiple comparisons. (C) Plasma IL-6 levels at the endpoints indicated in (H), Il-6 was below the assay detection limit for OT-KxPxCx^IL6^ mice. Statistically tested with 2-way ANOVA main effects only with Tukey correction for multiple comparisons. (A-C) N = 2 female, 3 male KxPxCx^IL6^, 1 female, 4 male KxPxCx^parental^ mice. (D) Representative pancreas cross sections at 5 (left) and 10 days (right) from OT-KxPxCx^parental^ (top) and OT-KxPxCx^IL6^ (bottom) tumor implantation. Scale bars represent 50 um (top) and 100 um (bottom). (E) Tumor area as a percent of total tissue area at 5 and 12 days, quantified by board-certified pathologist. (F) Percentage of tumor area classified as poorly-differentiated, infiltrative carcinoma at 5 days by board-certified pathologist. (G) Tumor mass at 5 and 12 days. (B-D) N = 3 male, 3 female mice per group. (H) Expression of *Il6 transgene* expression, measured by qPCR at 5 (N = 3 male, 3 female), 10 (N = 6 male), 17 (N = 3 male, 2 female), 24 (N = 3 male, 2 female), and 40 (N = 16 male, 16 female) days. Statistically tested with One-way ANOVA with Tukey correction for multiple comparisons. (I) Plasma IL-6 measured by ELISA at 5 (N = 10 male KxPxCx^Parental^, 11 male KxPxCx^IL6^), 12(N = 3 male, 3 female KxPxCx^Parental^, 3 male, 2 female KxPxCx^IL6^), and 17 (N = 2 male, 3 female KxPxCx^IL6^) days. Statistically tested with 2-way ANOVA main effects only with Tukey correction for multiple comparisons. (J) Body mass change as a percentage of initial body mass over time. N = 8 male, 8 female mice per group. Statistically tested with 3-way ANOVA. P<0.0001 for Time, Sex, Tumor Status, Time x Sex, Time x Tumor Status. p=0.0002 for Sex x Tumor Status. p= 0.2900 for Time x Sex x Tumor Status. (K) Gastrocnemius muscle mass normalized to initial body mass. (L) Atrophy-related gene expression (*Trim63* and *Fbxo32*) in gastrocnemius muscle, measured by qPCR. (J-L) 5d N = 1 female, 3 male mice per group. 10d N = 6 male PDAC, 5 male sham mice. Error bars represent SEM. Unless otherwise noted, 2x2 studies were statistically tested with a full effects model 2-way ANOVA and Sidak multiple comparisons test. 2-group analysis tested with unpaired t-test. **** P<0.0001, ***P<0.001, **P<0.01, *P<0.05.

OT-KxPxCx^parental^ plasma IL-6 levels were in line with previously published values, while OT-KxPxCx^IL6^ levels reached 100-fold higher levels at day 5, followed by undetectable levels at later time points (**Figure 1I**) ^10, 12^. Because high circulating IL-6 is associated with cachexia development in PDAC, we investigated total body and muscle mass^10^. High plasma IL-6 and peak tumor burden associated with decreased body mass, which recovered with the normalization of IL-6 levels after tumor clearance in the OT-KxPxCx^IL6^ model (**Figure 1J**). Gross muscle mass trended downward at five days and was significantly decreased at 10 days, while muscle atrophy gene (*Trim63* and *Fbxo32*) expression was elevated at 5 days and reduced to baseline at 10 days. These data reflect a delay in tissue recovery as the tumor resolves and body mass returns to baseline (**Figure 2K-L**). Collectively, these data show that IL-6 overexpression in PDAC cells leads to severe wasting, followed by body mass recovery upon spontaneous tumor resolution.

**Figure 2:**
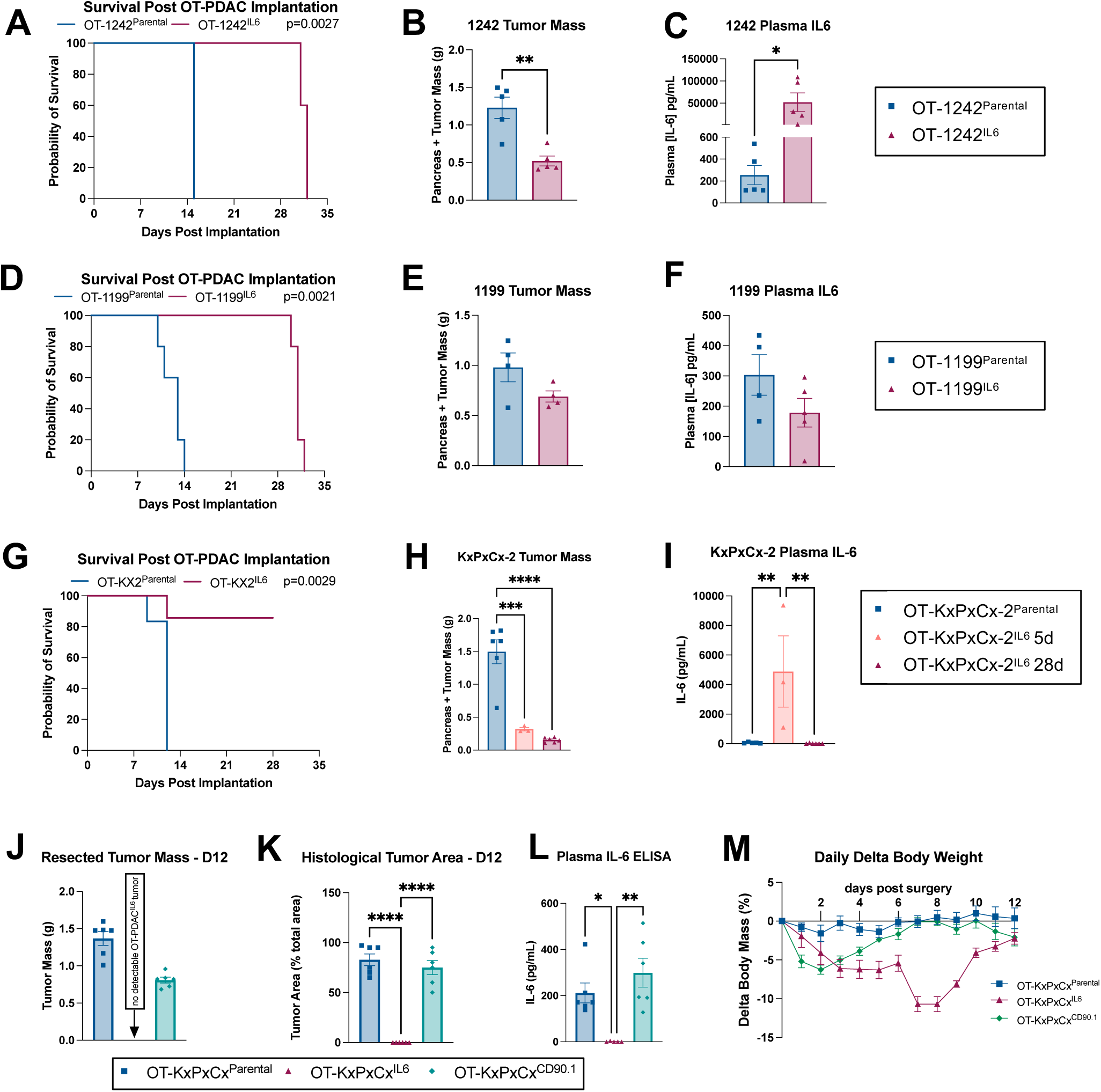
Tumor-cell IL-6 overexpression transduction controls. A) Survival comparison of Sham, OT-1242^Parental^ and OT-1242^IL6^. Statistically tested with Log-rank (Mantel-Cox) test on pairwise comparisons. (B) Pancreas and tumor mass at humane endpoint. (C) Plasma IL-6 levels at humane endpoint. (D) Survival comparison of Sham, OT-1199 ^Parental^ and OT-1199^IL6^. Statistically tested with Log-rank (Mantel-Cox) test on pairwise comparisons. (E) Pancreas and tumor mass at humane endpoint. (F) Plasma IL-6 levels at humane endpoint. (A-F) N= 5 male mice per group. (G) Survival comparison of Sham, OT-KX2^Parental^ and OT-KX2^IL6^. Statistically tested with Log-rank (Mantel-Cox) test on pairwise comparisons. (H) Pancreas and tumor mass at humane endpoint (OT-KX2^Parental^); 5 and 28 days (OT-KX2^IL6^). Statistically tested with one-way ANOVA. (I) Plasma IL-6 levels at humane endpoint. (G-I) N = 3 male mice OT-KX2^IL6^ d5, 6 male mice OT-KX2^IL6^ d28 and OT-KX2^parental^. (J-M) Control-transduced PDAC cells (OT-KxPxCx^CD90.1^) do not spontaneously clear. (J) Tumor mass at 12 days post implantation. (K) Tumor area as a percent of total tissue area at 12 days post implantation, quantified by board-certified pathologist. (L) Plasma IL-6 measured by ELISA at 12 days post implantation. (M) Body mass change as a percentage of initial body mass over time. Statistically tested with one-way ANOVA analysis with repeated measures. P<0.0001 for Time and Time x Tumor Status. p=0.0002 for Tumor Status. (J-M) N = 3 male, 3 female mice per group. Error bars represent SEM. Unless otherwise noted, 2-group studies were statistically tested with unpaired t-test. 3-group studies were statistically tested with 1-way ANOVA and Sidak multiple comparisons test. **** P<0.0001, ***P<0.001, **P<0.01, *P<0.05.

### Multiple OT-PDAC^IL6^ cell lines, but not CD90.1 transduction control lines, display improved survival outcomes

We then tested whether the impact of IL-6 overexpression on tumor clearance and mouse survival applied to other murine PDAC cell lines: KPC-1199^19^, and KPC-1242^20^. While both PDAC^IL6^ cell lines showed significantly improved survival, only OT-1242^IL6^ tumors remained smaller at humane endpoint, when compared to the respective parental control line (**Figure 2A-B, D-E**). High levels of IL-6 overexpression, quantified as plasma IL-6 levels at endpoint, correlated with less tumor burden: 1242^IL6^ caused ∼250 fold elevation of IL-6, and 1199^IL6^ did not show a significant overexpression or reduction in tumor mass (**Figure 2C,F**). We validated the clearance and survival phenotypes using an independently generated KxPxCx^IL6^ (KX2^IL6^) (**Figure 2G-I**). This led us to hypothesize that reduction in tumor mass was driven by IL-6 overexpression. We tested this by using a KxPxCx transduction control cell line, where we expressed CD90.1 alone without IL-6 (OT-KxPxCx^CD90.1)^. OT-KxPxCx^CD90.1^ mice replicated our findings with non-transduced OT-KxPxCx^parental^ mice. Both groups had reached humane euthanasia endpoint by 12 days, carcinoma covering approximately 80% of total tissue area, circulating IL-6 elevated to the expected 200-300 pg/mL range, and body mass loss not exceeding 10% (**Figure 2J-M, Table S1**). Finally, we tested the tumor response to IL-6 overexpression when fewer cells are implanted, thus prolonging the period of host exposure to tumor before humane euthanasia criteria are met. We tested 5,000 KxPxCx cells, 100,000 KPC-1199 cells, and 100,00 KPC-1242 cells, and found that survival, tumor mass, and plasma IL-6 replicated our findings with 1 million cell implantation doses (**Figure 3A-I**). We therefore concluded that the tumor clearance we observed was specific to IL-6 overexpression and most robust in the KxPxCx cell line, which we selected for continued testing. Because we did not observe a dose-dependent effect, we proceeded to exclusively test the implantation dose of 1 million KxPxCx^IL6^ cells, which aligns with our previously published work.^12^

**Figure 3:**
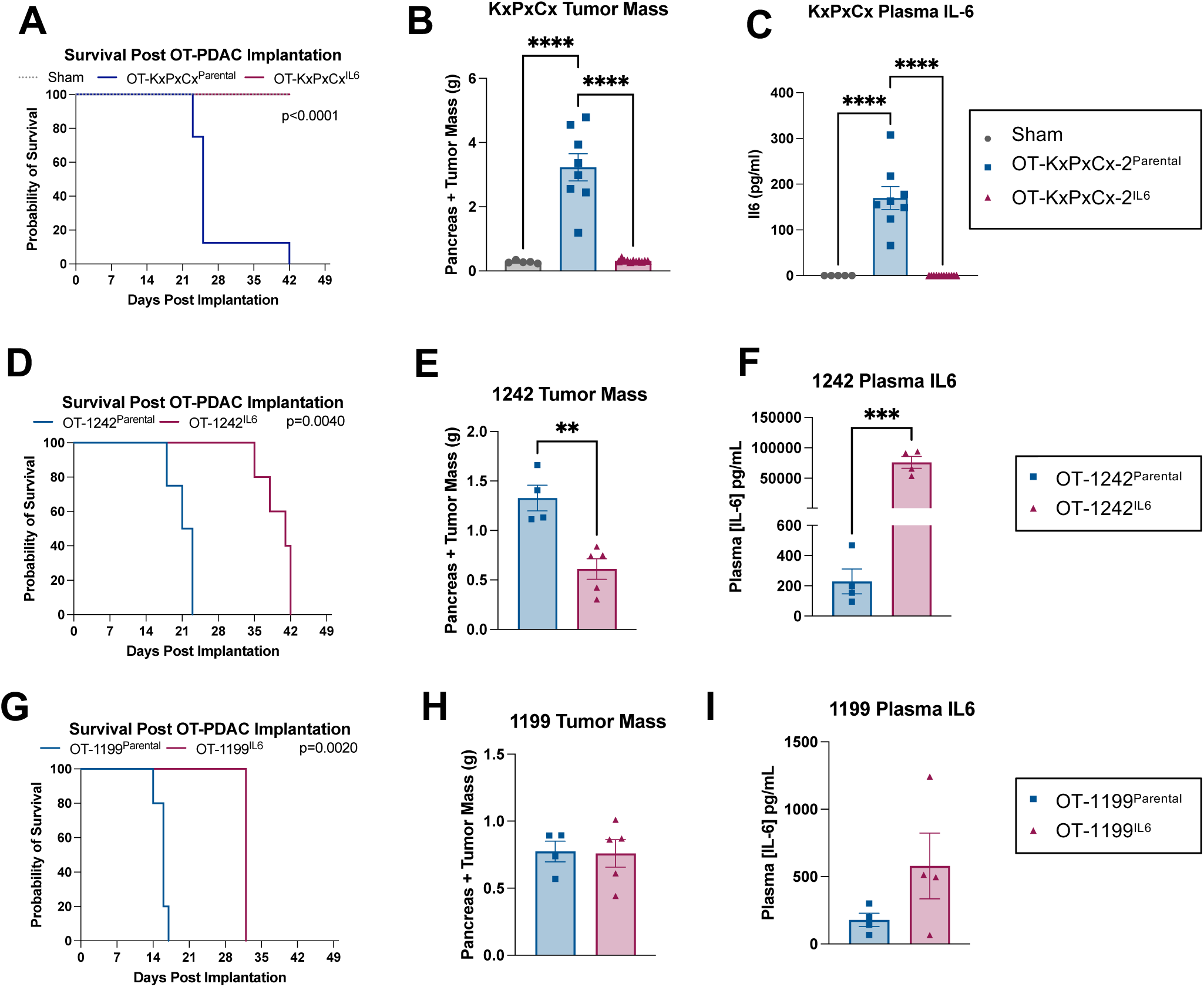
Effects of tumor-specific IL-6 overexpression are maintained with lower tumor inoculation. (A) Survival comparison of OT-KxPxCx^Parental^ and OT-KxPxCx^IL6^ mice implanted with 5,000 cells. Statistically tested with Log-rank (Mantel-Cox) test on pairwise comparisons. (B) Pancreas and tumor mass at humane endpoint (OT-KxPxCx^Parental^); 42 days (OT-KxPxCx^IL6^). (C) Plasma IL-6 levels at endpoint. (A-C) N = 11 male mice (OT-KxPxCx^IL6^), 8 male mice (OT-KxPxCx^Parental^), and 5 male mice (sham). (D) Survival comparison of Sham, OT-1242^Parental^ and OT-1242^IL6^. Statistically tested with Log-rank (Mantel-Cox) test on pairwise comparisons. (E) Pancreas and tumor mass at humane endpoint. (F) Plasma IL-6 levels at humane endpoint. (G) Survival comparison of Sham, OT-1199 ^Parental^ and OT-1199^IL6^. Statistically tested with Log-rank (Mantel-Cox) test on pairwise comparisons. (H) Pancreas and tumor mass at humane endpoint. (I) Plasma IL-6 levels at humane endpoint. Error bars represent SEM. Unless otherwise noted, 2-group studies were statistically tested with unpaired t-test. 3-group studies were statistically tested with 1-way ANOVA and Sidak multiple comparisons test. **** P<0.0001, ***P<0.001, **P<0.01, *P<0.05.

### Tumor IL-6 overexpression induces lymphocytic anti-tumor immune response

Spleens from OT-KxPxCx^IL6^ mice were significantly enlarged, consistent with IL-6-induced peripheral T cell expansion^15^ (**Figure 4A**). We therefore hypothesized that the extreme levels of IL-6 generated in the pancreas of OT-KxPxCx^IL6^ mice facilitated immune cell expansion and immune-mediated tumor clearance. We used flow cytometry to evaluate the immune profile of OT-KxPxCx^IL6^ and OT-KxPxCx^parental^ tumors and discovered that pancreata bearing OT-KxPxCx^IL6^ tumors were enriched with CD4^+^ T cells and Natural Killer (NK) cells, while Foxp3^+^ T-regulatory cells were decreased (**Figure 4B-I, S4-5**). We did not identify increases in Th17 cells, or Granzyme B^+^ CD8 T cells, which are two phenotypes associated with increased anti-tumor immunity and IL-6 signaling (**Figure S6-7**). We did, however, see a decreased ratio of Foxp3^+^ CD4 T cells to CD8 T cells, indicating a shift towards a less immunosuppressive tumor microenvironment (**Figure 4G**). Because increased tumor-infiltrating lymphocytes are associated with improved outcomes and therapeutic efficacy, we next assessed the localization of T cells relative to tumor cells^21^. By immunofluorescent staining, we found that OT-KxPxCx^parental^ and OT-KxPxCx^IL6^ mice had equal densities of CD3^+^ T cells in both tumor-associated stroma and tumor nests (**Figure 4J-L**). These data initially contradicted the flow cytometry data. However, upon histological assessment, we found that OT-KxPxCx^IL6^ pancreata exhibited a significantly greater number of tumor-associated lymphoid aggregates (**Figure 4M-N**). In PDAC, lymphoid aggregate formation is associated with improved survival^22, 23^. In OT-KxPxCx^IL6^ mice, the lymphoid aggregates were often near, but not necessarily located within the tumor stroma, accounting for the differences between our flow cytometry and immunofluorescence data. In summary, locally high IL-6 induces accumulation of lymphoid aggregates, increased CD4^+^ T cells, and decreased Foxp3^+^ T regulatory cells. These conditions favor the hypothesis that tumor clearance occurs via an anti-tumor T cell response.

**Figure 4:**
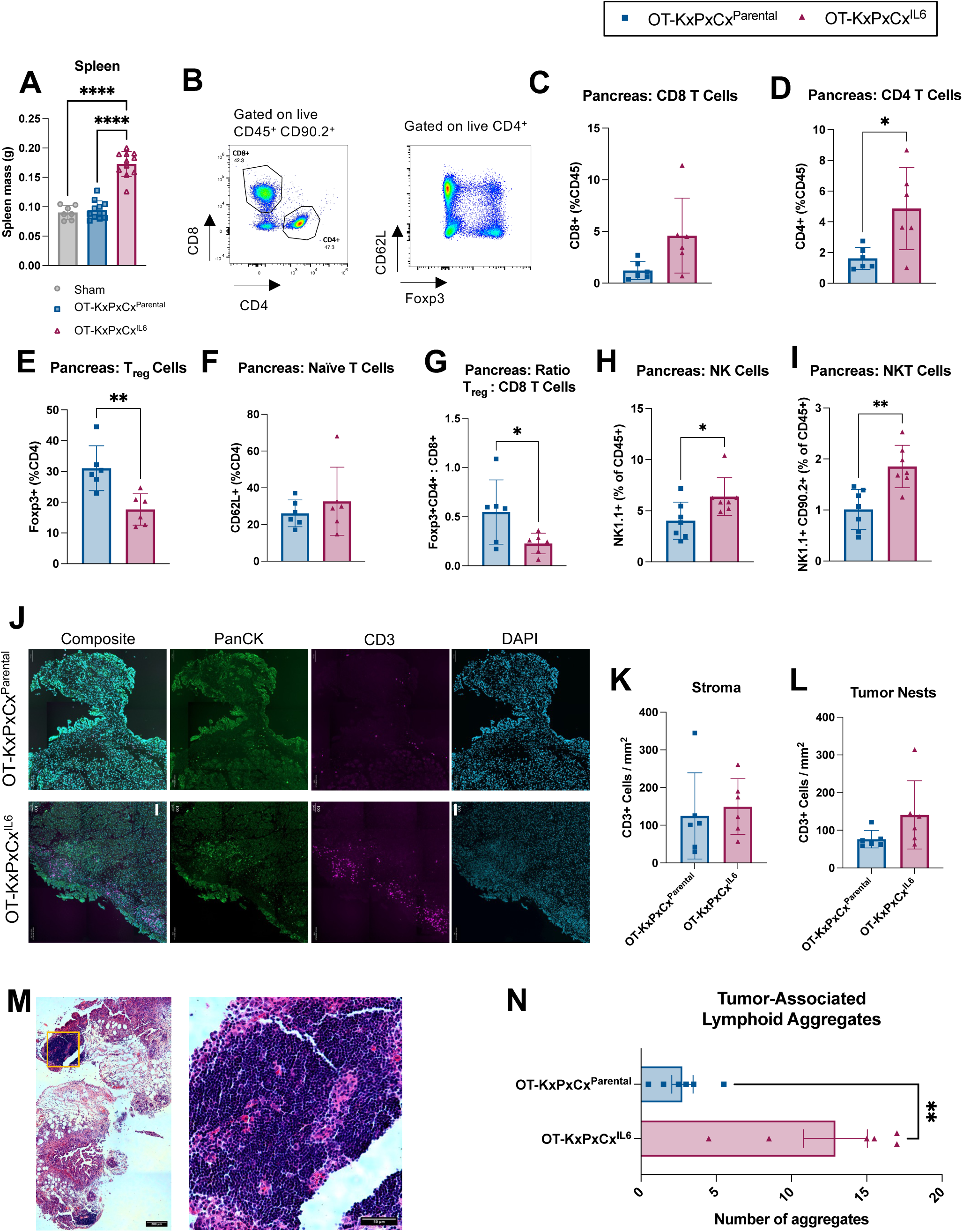
OT-KxPxCx^IL6^ induces lymphocytic anti-tumor immune response. (A) Spleen mass at 5 days. N = 6 male mice per group. (B-I) Intra-tumoral immune cell populations from pancreas at 5 days: (B) Representative flow gating strategy for T cells. (C) CD8^+^ T cells, (D) CD4^+^ T cells, (E) Foxp3^+^ T regulatory cells, (F) CD62L^+^ Naive T cells. (G) Ratio of Foxp3^+^CD4^+^ to CD8^+^ cells, representative of a the degree of anti-tumor activity in the tumor microenvironment. (H) NK1.1^+^ Natural Killer cells, (I) NK1.1^+^/CD90.2^+^ NK T cells. (B-F) N = 6 male mice per group. (G-I) N = 4 female, 3 male mice per group. (J) representative images of CD3^+^ T cell infiltration in tumor and associated stroma of OT-KxPxCx^parental^ (top) and OT-KxPxCx^IL6^ (bottom) mice. Tissues were stained with DAPI (blue), CD3 (magenta), and Pancytokeratin (PanCK, green). (scale bars = 100 um). (K-L) CD3^+^ cells counted per mm^2^ of stroma (K) or tumor nests (L) in immunofluorescence stained pancreas tissues. N = 3 male, 3 female mice per group. (M) Representative H&E image of lymphoid aggregates adjacent to tumor and stroma in OT-KxPxCx^IL6^ pancreas (left) and magnified image of lymphoid aggregate (right). Area of magnification is denoted on left image with orange box. (N) Pathologist-evaluated number of tumor-associated lymphoid aggregates in H&E stained pancreas cross sections from tissue collected at 5 days. N = 3 male, 3 female mice per group. Error bars represent SEM. 2-group analysis tested with unpaired t-test. **** P<0.0001, ***P<0.001, **P<0.01, *P<0.05.

### T cells are necessary for OT-KxPxCx^IL6^ tumor clearance

We hypothesized that T cells are necessary for OT-KxPxCx^IL6^ tumor clearance, which we tested via antibody-based CD4 and CD8 depletion in tumor-bearing mice, compared to IgG control treatment. Mice received antibody injections two days before tumor implantation and every four days following. We validated T cell depletion via flow cytometry for total T cells (CD90.2), and CD4^+^ and CD8^+^ cells (**Figure 5A-C**). We followed humane endpoints for the CD4/CD8-depleted group, which reached euthanasia criteria at 8-11 days post-implantation (**Figure 5D**). We euthanized all sham and IgG-treated mice when all CD4/CD8-depleted mice were euthanized, although they were healthy at the time. We used a second engineered cell line, KxPxCx^IL6-LUC^, which co-expressed IL-6 and luciferase, to measure tumor burden over time. Both OT-KxPxCx^IL6-LUC^ longitudinal data, and OT-KxPxCx^IL6^ endpoint tumor mass data show that CD4/CD8-depletion prevents tumor clearance (**Figure 5E-G**). Histological evaluation by a board-certified pathologist found tumor present in 10/10 CD4/CD8-depleted mice, and in 1/8 IgG control mice (**Figure 5H**). This was supported by *Il6*-transgene qPCR data and plasma IL-6 levels, which showed no elevation of IL-6 in 6/8 IgG-treated PDAC mice (**Figure 5I-J**). We conclude that CD4 and CD8 T cells are necessary for OT-KxPxCx^IL6^ tumor clearance.

**Figure 5:**
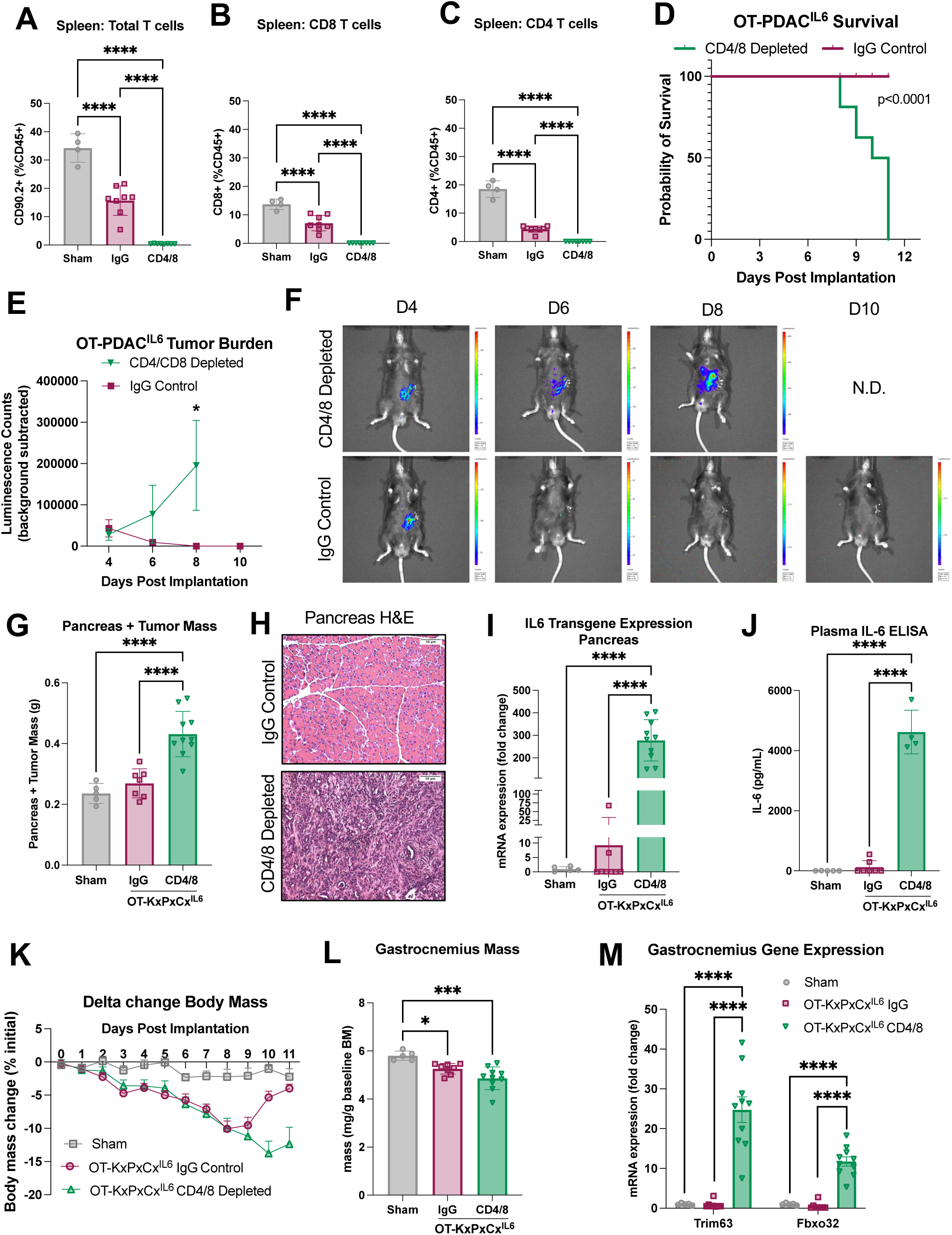
CD4^+^ and CD8^+^ T cells are necessary for OT-KxPxCx^IL6^ tumor clearance. (A-C) Confirmation of CD4/CD8a T cell depletion via flow cytometry on spleen 12 days post implantation for (A)CD90.2+ total T cells, (B) CD8+ T cells, (C) CD4+ T cells. (D) Survival comparison of OT-KxPxCx^IL6^ mice given CD4/CD8 depletion or IgG control antibodies. Statistically tested with Log-rank (Mantel-Cox) test. (E) OT-KxPxCx^IL6-LUC^ tumor growth, measured by IVIS imaging. Statistically tested with 2-way ANOVA, for time points with representation from both groups, with Šídák multiple testing correction. (F) Representative images from IVIS imaging of OT-KxPxCx^IL6-LUC^ 10 minutes after luciferin injection. Images are of the same animals longitudinally. CD4/CD8 depleted mice reached humane endpoint prior to D10 scan. (G) Pancreas and tumor mass at endpoint. (H) Representative pancreas cross sections at humane euthanasia endpoint from IgG control (top) and CD4/CD8 depleted (bottom) mice. Scale bars represent 100 um. (I) expression of *Il6 transgene* expression, measured by qPCR. (J) Plasma IL-6 measured by ELISA. (K) Body mass change as a percentage of initial body mass over time. Statistically tested with Mixed-effects analysis with repeated measures. P<0.0001 for Time and Time x Tumor Status. p=0.0048 for Tumor Status. (L) Gastrocnemius muscle mass at humane euthanasia endpoint normalized to initial body mass. (M) Atrophy-related gene expression (*Trim63* and *Fbxo32*) in gastrocnemius muscle, measured by qPCR. All male mice, N = 5 sham, 10 KxPxCx^IL6^ per antibody treatment group. Error bars represent SEM. 3 group studies were statistically tested with a 1-way ANOVA and Tukey correction for multiple comparisons. **** P<0.0001, ***P<0.001, **P<0.01, *P<0.05.

Because we previously saw recovery of wasting phenotypes in OT-KxPxCx^IL6^ mice as the tumor resolved, we also investigated cachexia resolution in T cell-depleted mice. CD4/CD8 depletion prevented body mass recovery and led to sustained muscle wasting as evidenced by decreased muscle mass and increased atrophy-related gene expression (*Trim63, Fbxo32*) (**Figure 5K-M**). The recovery in wasting is therefore directly associated with immune-mediated tumor clearance.

### OT-KxPxCx^IL6^ induces a durable T cell response to OT-KxPxCx^Parental^ tumors

Given the dependency of the anti-OT-KxPxCx^IL6^ response on T cells, which are known to elicit immunologic memory, we hypothesized that OT-KxPxCx^IL6^ tumors would generate a durable immune response that would protect mice in the case of a second tumor exposure. We first tested this hypothesis in a cohort of mice that had recovered from sham surgery or OT-KxPxCx^IL6^ for over two months, and were then rechallenged with OT-KxPxCx^parental-LUC^ (luciferase-expressing) at 76 days after the initial surgery (**Figure 6A**). Sham-recovered mice implanted with KxPxCx^parental-LUC^ reached euthanasia criteria 13-14 days after rechallenge implantation. In contrast, there were no deaths following rechallenge in the OT-KxPxCx^IL6^-recovered group (**Figure 6B**). Tumor burden measured longitudinally by IVIS and terminally (in sham-recovered mice only) revealed significant tumor growth only in sham-recovered mice (**Figure 6C-D**). Sham-recovered mice also lost more body mass than OT-KxPxCx^IL6^-recovered mice following rechallenge (**Figure 6E**). These data indicate that the potent anti-tumor immune response elicited by high concentrations of IL-6 is durable and, once established, not dependent on coincident supraphysiologic IL-6 levels. We then used CD4/CD8 antibody depletion to determine whether the tumor clearance during rechallenge was indeed T cell mediated. In this study, all mice recovered from OT-KxPxCx^IL6^ for 26 days before starting antibody treatment and were given sham surgery or implanted with OT-KxPxCx^parental^ tumors on day 28. We monitored mice for 12 days, until CD4/CD8-depleted mice reached humane euthanasia criteria (**Figure 6F**). We confirmed that clearance of the rechallenge OT-KxPxCx^parental^ tumor is dependent on T cells, which were effectively depleted by the antibody treatment (**Figure 6G-I**). Furthermore, we confirmed that CD4/CD8-depletion does not result in the outgrowth of potentially covert KxPxCx^IL6^ cells, as no tumor was present in any of the OT-KxPxCx^IL6^-recovered, sham-rechallenged, CD4/CD8-depleted mice (**Figure 6G**). These data show that the T cell response induced by OT-KxPxCx^IL6^ provides durable protection against molecularly similar PDAC tumor growth.

**Figure 6:**
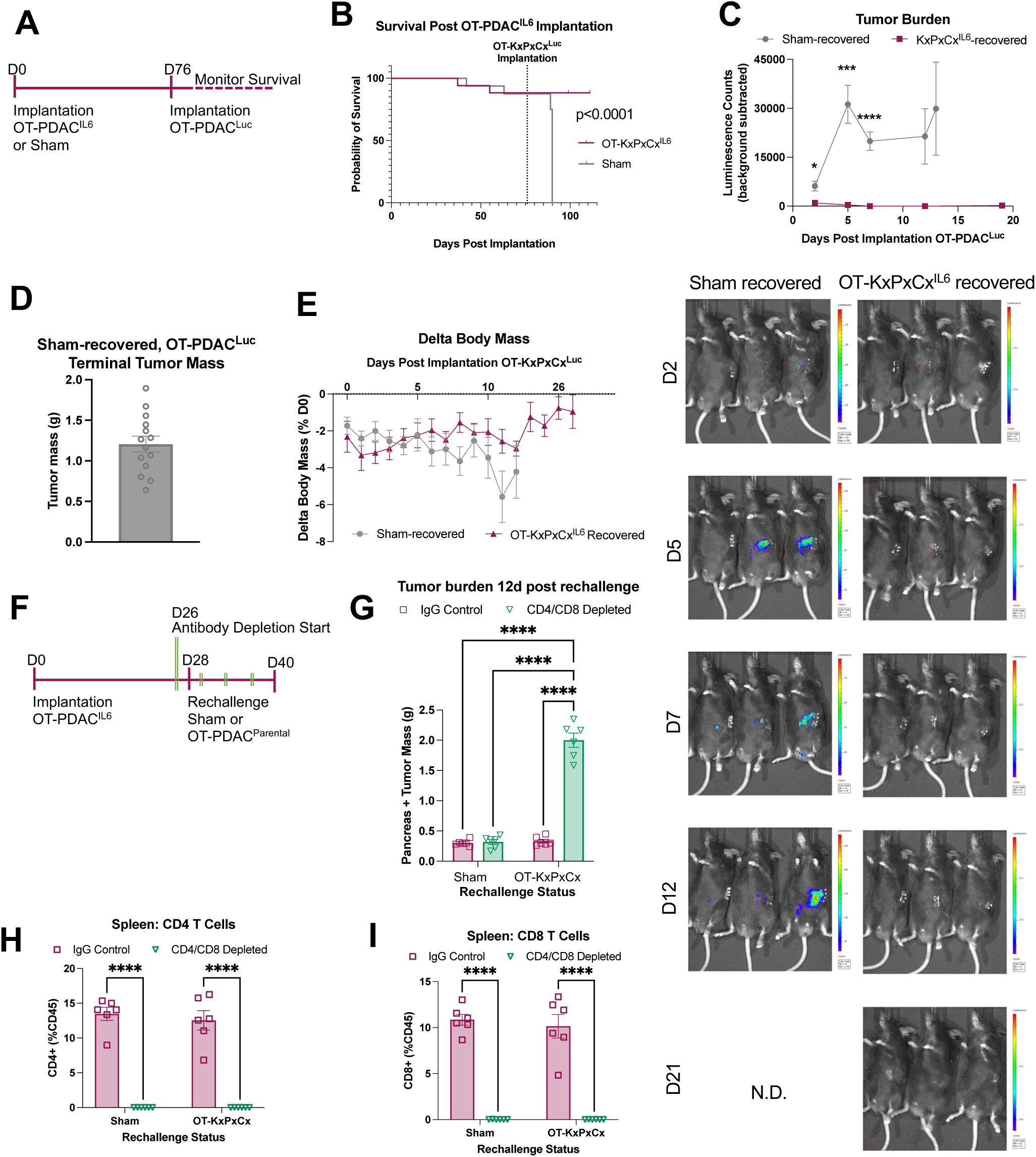
OT-KxPxCx^IL6^ induces a durable T cell response to OT-KxPxCx^Parental^ tumors. (A) Schematic timeline for B-E. Mice were implanted with OT-KxPxCx^IL6^ or given sham surgery, then all mice were rechallenged with OT-KxPxCx^Parental-LUC^ after 76 days. (B) Survival comparison of OT-KxPxCx^IL6^-recovered and sham-recovered mice. Statistically tested with Log-rank (Mantel-Cox) test. (C) OT-KxPxCx^Parental-LUC^ tumor growth, measured by IVIS imaging. Statistically tested with mixed effects model and Šídák multiple testing correction, using imputation to match KxPxCx^IL6^ endpoint with the sham endpoint. Corresponding IVIS images below. (D) Tumor mass at humane euthanasia endpoint for sham-recovered OT-KxPxCx^Parental-LUC^ tumors. (E) Body mass change as a percentage of initial body mass over time. Statistically tested with mixed effects model and Šídák multiple testing correction at timepoints where both groups were represented. p=0.0435 for Time, p=0.4903 for Tumor Status, p=0.0002 for Time x Tumor Status. (A-E) N = 7 male KxPxCx^IL6^, 8 male sham, 7 female KxPxCx^IL6^, 6 female sham. (F) Schematic timeline for G-I. Mice were implanted with OT-KxPxCx^IL6^ or given Sham surgery. 28 days later, mice were rechallenged with OT-KxPxCx^Parental^, and given CD4/CD8 depletion or IgG control antibodies beginning on D26 and every 4 days thereafter (denoted with double green lines). (G) Pancreas/tumor mass at 12 days. (H) Intra-tumoral CD4^+^ T cells at 12 days. (I) Intra-tumoral CD8^+^ T cells at 12 days. (F-I) All male mice N = 6 per group. Error bars represent SEM. 2x2 studies were statistically tested with a full effects model 2-way ANOVA and Sidak multiple comparisons test. **** P<0.0001, ***P<0.001, **P<0.01, *P<0.05.

### Lipid nanoparticle delivery of IL-6 mRNA replicates the anti-tumor effects observed in OT-PDAC^IL6^

To test whether the anti-tumor effects of elevated IL-6 in the tumor microenvironment could be derived through a therapeutically-relevant method, we engineered lipid nanoparticles (LNPs) that deliver IL-6 mRNA, are administered by IP injection, and target the pancreas. We used SM-102 LNPs containing luciferase mRNA to validate the particle delivery 6 hours after IP injection in healthy mice (**Figure 7A**). We found that luciferase expression was highest in the pancreas, but was also expressed in the liver, and was nearly undetectable in the spleen (**Figure 7A-B**). Following validation of dosing, we designed an experimental setup with escalating doses of LNPs, which were administered 3 hours prior to OT-PDAC^parental^ (no IL-6 overexpression by the tumor) implantation and every other day after (**Figure 7C**). We included an empty LNP group that was dose-matched for LNP cholesterol content to control for potentially immunogenic effects of the LNPs alone. We found that OT-PDAC^parental^ mice injected with IL-6 LNPs lost significant body mass and had approximately 50-fold higher circulating IL-6 than empty LNP or no LNP control OT-PDAC^parental^ mice (**Figure 7F-G**). IL-6 LNP OT-PDAC^parental^ mice euthanized 8 days post-implantation had significantly lower tumor burden and larger spleens (**Figure 7G-H**). We used flow cytometry to evaluate the tumor immune profile and found that tumors from OT-PDAC^parental^ mice injected with IL-6 LNPs did not have significantly more CD4^+^ or CD8^+^ T cells (**Figure 7I-J, S9**). Despite this, there were significantly fewer T-regulatory cells within the CD4 population, and a decreased ratio of Foxp3^+^ CD4 T cells to CD8 T cells in IL-6 LNP-treated OT-PDAC^parental^ mice, replicating our findings in OT-PDAC^IL6^ (**Figure 7K-L**). Finally, we saw significant enrichment of NK and NK T cells in the pancreas of IL-6 LNP-treated OT-PDAC^parental^ mice (**Figure 7M-N**). There were no differences in immune populations between the empty LNP and no LNP groups, suggesting that, in the pancreas with this dosing paradigm, empty SM-102 LNPs did not induce a T cell response. Together, these data show that IL-6 mRNA delivery to the pancreas results in an anti-tumor immune landscape and lower tumor burden in OT-PDAC^parental^ mice.

**Figure 7:**
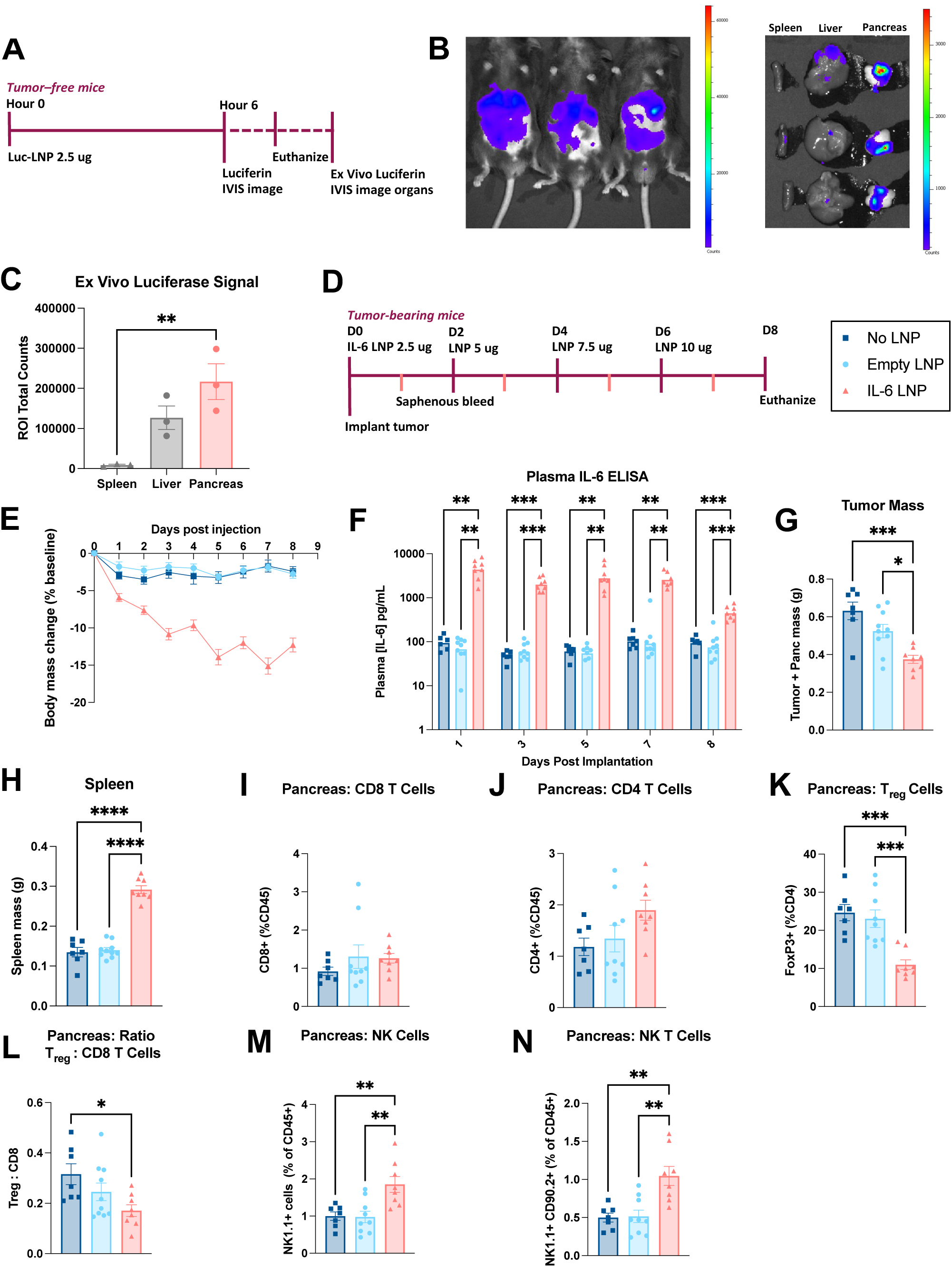
Lipid nanoparticle delivery of IL-6 mRNA replicates the anti-tumor effects observed in OT-PDAC^IL6^. (A) Schematic of luciferase LNP validation. Mice were injected with LNPs containing a total of 2.5 ug luciferase mRNA 6 hours prior to IVIS imaging, euthanasia, tissue collection, and tissue IVIS imaging. N = 3 male mice per group. (B) left- IVIS images of mice injected intraperitoneally with LNPs containing luciferase. Right – IVIS image of the spleen, liver, and pancreas, incubated ex vivo with luciferin prior to imaging. (C) Comparison of total luminescence counts in the pancreas and liver. Statistically tested with one-way ANOVA and Tukey multiple testing correction. (D) Schematic timeline for study presented in D-N. Mice were injected with LNP dose 1, 3 hours prior to tumor implantation. Mice were then injected with subsequent doses on 2, 4, and 6 days post-implantation. Blood was collected by saphenous bleed 1, 3, 5, and 7 days post-implantation. Mice were euthanized on day 8. N = 7 (No LNP), 10 (empty LNP), 8 (IL6 LNP). All mice were male. Error bars represent SEM. (E) Body mass change as a percentage of initial body mass over time. Statistically tested with two-way ANOVA P<0.0001 for Time, Treatment, and Time x Treatment. (F) Plasma IL-6 measured by ELISA at 1, 3, 5, and 7 days post-implantation, and at euthanasia (D8). Statistically tested with repeated measures mixed-effects model with Tukey correction for multiple testing. (G) Tumor and pancreas mass. (H) Spleen mass. (I-N) Intra-tumoral immune cell populations from pancreas: (I) CD8^+^ T cells, (J) CD4^+^ T cells, (K) Foxp3^+^ T regulatory cells, (L) ratio of Foxp3^+^ to CD8^+^ cells, representative of a the degree of anti-tumor activity in the tumor microenvironment, (M) NK1.1+ NK cells, (N) NK1.1+/CD90.2+ NK T cells. 3-group analysis tested with one-way ANOVA with Tukey correction for multiple testing. **** P<0.0001, ***P<0.001, **P<0.01, *P<0.05.

## DISCUSSION

Our model of PDAC IL-6 overexpression induced a robust, rapid, and durable anti-tumor T cell response, accompanied by rapid and severe wasting that recovered as the tumor was cleared. Although IL-6 is traditionally viewed as a negative factor in pancreatic cancer ^8–11^, we provide evidence that supraphysiologic levels of IL-6, achieved through tumor cell overexpression of IL-6 protein or LNP delivery of IL-6 mRNA, were sufficient to induce an anti-tumor immune landscape in the pancreas. The OT-PDAC^IL6^ tumor microenvironment was characterized by increased lymphoid aggregate formation, elevated CD4^+^ T cells, decreased Foxp3^+^ Treg cells, and elevated NK cells. LNP delivery of IL-6 also drove decreased Treg cell and increased NK cell infiltration. While the KxPxCx tumor line implanted orthotopically exhibited complete, recurrence-free survival, subcutaneous tumors of the same line displayed less complete and slower regression in tumor size. IL-6 overexpression in other KPC cell lines (1199 and 1242) resulted in significant improvements in mouse survival time, although tumors were never fully cleared, and mice progressed to humane euthanasia endpoints. These cases highlighted that the magnitude of anti-tumor immunity due to IL-6 overexpression was partially dependent on cell-line intrinsic properties.

Unlike cancer types that are now successfully treated with immunotherapy, survival rates for PDAC patients have increased very slowly over the past decade^24^. PDAC is highly immunosuppressive, causing immunotherapies, such as checkpoint blockade, to be ineffective clinically ^25^. Previous work indicated that enhancing T cell activation using exogenous agents, such as agonistic anti-CD40 antibody, improved response to checkpoint blockade therapies and PDAC tumor regression^1^. The Lesinski Group also used IL-6 blockade to enhance the effects of PD-L1 or CTLA4 blockade. Combining checkpoint inhibitors with anti-IL-6 reduced tumor growth and improved T-cell infiltration in implanted murine PDAC models^26, 27^. However, anti-IL-6 was only effective in bolstering the anti-tumor effects of immune checkpoint blockade; the IL-6 antibody did not exert anti-tumor effects on its own. Our work raises a fundamental contradiction regarding the role of IL-6 in PDAC and provides a basis for enhanced IL-6 expression as an alternative method to improve T cell response.

We propose that the effect of IL-6 on PDAC growth is pleiotropic and concentration-dependent. At low concentrations, IL-6 aids tumor development via signaling directly on neoplastic cells to drive transformation and growth. This is evidenced by the reduced frequency of high-grade pancreatic intraepithelial neoplasia observed in an aggressive genetic model of murine PDAC, when mice were treated with combination PD-L1 and IL-6 blockade^26^. Although some murine PDAC cells produce IL-6, prior work showed that the source of IL-6 in human PDAC tumors and orthotopic mouse models is primarily fibroblasts and immune cells^26, 28^. Combination anti-IL-6 and anti-PD-L1 therapy reduced the number of αSMA+ stromal cells, the loss of which potentially contributed to limiting PDAC progression in that model^26^. Conversely, at supraphysiologic concentrations seen in two independent methods of concentrated IL-6 expression in the pancreas (OT-KxPxCx^IL6^ and IL-6 LNP treatment), IL-6 stimulated an anti-tumor immune response. Circulating levels of IL-6 were approximately 100 times higher in OT-KxPxCx^IL6^ and 50 times higher after IL-6 LNP treatment than what we detected in OT-KxPxCx^parental^ mice. We presume that local concentrations in the pancreas were even higher in both systems. We can infer that pancreatic IL-6 is the initiating signal for T cell accumulation and eventual tumor clearance; however, the precise manner in which IL-6 mediates this remains unknown. In addition to increased numbers of tumor-infiltrating T cells, we detected increased NK cells, and decreased Treg cells intratumorally. It is likely that IL-6, which is a known immunomodulatory cytokine, impacted multiple cell populations simultaneously to orchestrate an anti-tumor immune microenvironment^29^. Conversely, it is also likely that lower levels of IL-6 promote expansion of stromal cells, which can contribute to an immune-deplete tumor microenvironment. Future work might investigate how high levels of IL-6, as seen in our OT-KxPxCx^IL6^ model, influence stromal development in PDAC tumors.

IL-6 was also established as a driver of acute-phase protein response in the liver, which primed the liver for increased metastasis formation^30^. The KxPxCx cell line used most extensively in this study is not highly metastatic when implanted orthotopically. Liver metastasis was only detected in one mouse at 5 days post-implantation in the OT-KxPxCx^IL6^. At 12 days post-implantation, zero mice had liver metastases detectable by qPCR (**Figure S2**). Given the high circulating levels of IL-6 in our models, it is likely that STAT3 was activated in hepatocytes, which could contribute to the rapid onset of cachexia in OT-PDAC^IL6^ mice^12^. However, we did not directly assess the activation of acute-phase protein response in our model. The rapid resolution of the OT-PDAC^IL6^ primary tumor and micro-metastatic disease made this particular model incompatible with rigorously assessing how supraphysiologic IL-6 affected the hepatic metastatic niche. It would be necessary to further investigate the impact of IL-6 on hepatic inflammation if IL-6 elevation were pursued therapeutically in the future, to avoid potentially unexpected negative consequences for metastatic disease.

In this work, we confirmed the anti-tumor effects of IL-6 overexpression in PDAC cells using IL-6 LNPs that targeted the pancreas. LNPs are rising in interest as therapeutic vectors for mRNA delivery due to their clinical safety and ability to customize particle content and delivery method to improve tissue-specific targeting^31–34^. Tissue specificity is especially attractive for anti-tumor therapeutic delivery that minimizes off-target effects^35, 36^. Recent publications show that next-generation ionizable lipids can improve pancreas specificity of LNPs^37, 38^. LNP-delivered IL-12 was sufficient to prevent PDAC growth^37, 38^. This, and our own work with IL-6, adds to a growing body of literature showing that local expression of immunomodulatory cytokines leads to an anti-tumor T cell response^5–7^. Further work is needed to understand how the T cell response to cytokine overexpression might be conserved and how it could be best translated to clinical use.

Our data support the widely-accepted notion that IL-6 induces acutely negative effects, as evidenced by rapid body mass loss of 10-15% in OT-KxPxCx^IL6^ mice and mice that received IL-6 LNPs (**Figure 2H**). In OT-KxPxCx^IL6^ mice, which derived IL-6 expression primarily from the tumor cells, tumor clearance led to rapid recovery in body mass. For patients with PDAC, who are already vulnerable to weight loss, the pro-cachectic effects of IL-6 would likely limit the clinical utility of therapies that elevate IL-6 systemically. Because IL-6 is a secreted protein, it is likely that even pancreas-directed delivery, for example targeted LNP delivery of mRNA, would pose a high risk of side effects for patients. Instead, future work will focus on identifying the immune subpopulations that integrate IL-6 signaling into a T cell response, with the goal of identifying targetable drivers of the anti-tumor response. Using IL-6 overexpression as a model of effective PDAC clearance will open doors to understanding IL-6-mediated T cell activation in PDAC and the intricacies of cachexia resolution after PDAC tumor clearance.

## METHODS

### Mouse Studies

#### Husbandry

C57BL/6J (WT, JAX 000664) mice were purchased from The Jackson Laboratory (Bar Harbor, ME) and maintained in our animal facility. All mice were housed and bred in a dedicated mouse room maintained at 26 °C, 40% humidity, and 12-h light/dark cycle. Mice were provided *ad libitum* access to food and water (5L0D, PicoLab) unless otherwise stated. All mice were 12 weeks of age at experiment start. Sex in each experiment is defined in the figure legends. When single housed, mice were allowed a 7-day acclimation period prior to procedure/study start. All tumor studies followed humane endpoints. All mice were humanely euthanized via cardiac puncture or cervical dislocation under deep isoflurane anesthesia. Mouse studies were conducted in accordance with the NIH Guide for the Care and Use of Laboratory animals, and approved by the Oregon Health & Science University IACUC.

#### Orthotopic Tumor Implantation

A vial of frozen KPC cells was thawed prior to each implantation, and 1 million cells were implanted in 23 uL of PBS per mouse. All mice were anesthetized with isoflurane, scrubbed with betadine, and a para-midline incision was made in the abdomen to expose pancreas. KPC cells or vehicle (PBS) were injected directly into the pancreatic parenchyma. Pancreas was placed back into position and incision was closed using two sutures (4-0 Polysorb) and two skin staples.

#### Subcutaneous Tumor Implantation

A vial of frozen KPC cells was thawed prior to each implantation, and 1 or 2 million cells were implanted in 100 uL of PBS per mouse. The mouse’s lower right abdomen was shaved, then the needle was inserted near the right 4^th^ mammary gland.

#### Antibody-based depletion

Mice were dosed intraperitoneally (IP) with either a combination of CD4 (BioXcell, #BE0003-1) and CD8a (BioXcell, #BE0061) depletion antibodies, or Rat IgG2b isotype control antibody (BioXcell, #BE0090), which were resuspended in InVivoPure ph7.0 Dilution Buffer (BioXcell, #IP0070) per the manufacturer’s instructions. First dose was 0.2 mg each antibody per mouse, given IP two days prior to tumor implantation. Following doses were 0.1 mg each antibody per mouse, given IP every 4 days after initial dose.

#### Lipid nanoparticle administration

LNPs were injected with a 30g insulin syringe intraperitoneally, following the dosing scheme outlined in Figure 7A and D. LNPs were dosed based off of RNA content and were diluted with calcium- and magnesium-free PBS. Empty LNP dose was calculated by matching the cholesterol content in the empty and IL-6-containing LNPs.

#### Saphenous blood collection

We collected blood via saphenous bleeds during the LNP study. Mice were restrained by scruffing while occluding the vein, and the hindlimb was wiped with petroleum jelly to expose the vein. A 23G needle was used to gently prick the saphenous vein. Blood was collected in EDTA-treated tubes, which were then placed on ice. Bleeding was stopped with pressure and mice were returned to the cage. Each bleed alternated between left and right hindlimb. Approximately 40 uL of blood was collected at each timepoint. Samples were spun down for 20 minutes at 2,000xg and plasma was stored at -80C.

### Cell lines

#### Growth Conditions and validation

All cells were maintained at 37°C and 5% CO_2_ in a humidified incubator, and tested negative in house for mycoplasma using Universal Mycoplasma Detection Kit (30-1012K). *Kras^G12D/+^, Tp53^R172H/+,^ Pdx1-Cre* (KPC) cell lines was generously shared by Dr. Elizabeth Jaffee (KxPxCx) and Dr. David Tuveson (1199 and 1242)^18–20, 39^. KPC cells were grown on tissue culture-treated dishes in growth media consisting of RPMI (Gibco) with 10% FBS (Corning) and 1% penicillin/streptomycin (Gibco).

#### Engineered KPC

KPC cells expressing the surface marker Thy1.1 (CD90.1) with blasticidin resistance (BSR), IL-6 with puromycin resistance, and luciferase with hygromycin resistance, were generated from our stock of KPC cells (female) described above. IL-6 sequence was codon-optimized for efficient expression (**Figure S9**). Platinum-E ecotropic packaging cells were transfected with plasmid DNA encoding MSGV-Thy1.1, MSGV-IL6-Puro, or MSGV-Luciferase as described previously^40^. Retroviral supernatants were spiked with 2ug/mL polybrene and were mixed 1:1 with fresh media before adding to 6-well tissue culture treated plates. Cells were spun at 2000g for 90min, 32C, no brake. Cells were then incubated at 37C for 48 hours before washing off the viral supernatant and adding DMEM media (Gibco) supplemented with 10% FBS (Corning). Two days later, KPC cells were placed in complete DMEM media containing puromycin (5ug/mL) and/or blasticidin (5ug/mL) and/or hygromycin (500 ug/mL) to select for transduced cells. Following antibiotic selection, successful transduction was confirmed via flow cytometry staining for Thy1.1. KxPxCx^CD90.1^, KxPxCx^IL6^, and KxPxCx^LUC^ cells were implanted for OT-PDAC as described for parental KPC cells. Continued culturing of engineered cells was done in selection media described.

### Flow cytometry

*Brefeldin A injections:* For intracellular cytokine staining for flow cytometry, we followed previously published protocols for golgi transport blockade^41^. Briefly, each mouse received 100ug Brefeldin A (Selleckchem) injected retro-orbitally 5 hours prior to tissue collection.

#### Sample preparation

We collected tumors from mice 5 days post implantation, and tumors were weighed, then placed in PBS on ice. Tumor tissue was minced and digested (7). After dissociation, we strained tumor suspension through at 100 um filter, and performed ACK lysis. We collected spleens from mice at the endpoint specified. Spleens were pressed through a 70 um filter, rinsed with PBS, pelleted at 1500 RPM for 5 minutes, and lysed with ACK lysis.

#### Staining

We stained samples with live/dead stain (1:2000) and surface protein antibodies (1:200 each), and incubated for 20 minutes room temperature (**Table S3**). After staining, we washed samples with FACS buffer and pelleted. For intracellular staining (Foxp3), we fixed and permeabilized cells with 4% paraformaldehyde (BD Cytofix/Cytoperm), washed cells, and then resuspended cells in antibody diluted 1:200 in permeabilization buffer (BD Perm/Wash) with overnight incubation at 4C . The next day we washed and resuspended cells with FACS buffer prior to analysis.

#### Instrumentation and analysis

All samples were analyzed in the OHSU Flow Cytometry Shared Resource using the Cytek Aurora flow cytometer (Cytek Biosystems), data was analyzed in FlowJoTM v10.8.1. Samples were first gated according to size and single cells, then all live cells were captured. The live population was gated on CD45+ cells to capture all leukocytes. To identify T cells, leukocytes were gated on CD90.2 then sub-gated for CD4+ or CD8+ cells. The CD4+ T cell population was gated on Foxp3 to assess T regulatory cells. Natural killer (NK) cells were sub-gated from the parental CD45+ gate. All NK1.1+ cells were captured, and then divided by expression of CD90.2 to classify as NK (conventional, CD90.2-) or NK T cells (CD90.2+).

### Histology

Pancreas/tumor tissue was fixed overnight in 4% PFA, then stored in 70% ethanol. Tissues were paraffin embedded, sectioned, and hematoxylin and eosin (H&E) stained by the OHSU Histopathology Shared Resource. Tumor tissue was sectioned in 5um slices at levels 50 um apart. H&E stained tumor cross-sections were evaluated by a board-certified gastrointestinal pathologist. All samples were blinded during sectioning/staining and during evaluation. 3-5 depths of tissue were qualitatively assessed per mouse. For day 5 tumor samples, 2 depths of tissue were quantified per mouse and averaged together. For day 12 tumor samples, 1 section was quantified.

### Immunofluorescent staining and quantification

Pancreas/tumor tissue was dissected, transferred to BD Cytofix/Cytoperm diluted to 1% PFA, and kept at 4C overnight. Tissue was then transferred to 30% sucrose in PBS and kept at 4C overnight. Tissue was then washed twice in PBS before embedding in OCT media (Sakura). Tissue was cut at 8 um onto superfrost plus slides and stored at -80C until staining. To stain, slides were washed with PBS, blocked with 2.5% BSA 0.3% TritonX in PBS for one hour, stained with pre-conjugated antibodies (**Table S3**) for 1h, washed with PBS, quenched with TrueView Autofluorescence Quench kit (Vector, SP-8400) 2-5 min, stained with DAPI diluted 1:1000 for 10 min, and mounted with Vectashield Vibrance mounting medium (Vector, H-1700). Whole tissue sections were imaged on a Zeiss Axio Scan 7 at 20x magnification. Two tissue sections at least 344 um apart were assessed per sample. All images were blinded prior to annotation and analysis. In QuPath (V0.5.1), areas of tumor, as defined as PanCK positive, and stroma, as defined as PanCK negative abnormal tissue, were annotated. Adjacent tissue sections were stained with H&E and used as reference for areas of stroma. Annotations were made using only the DAPI and PanCK stains. After annotation, CD3^+^ cells (CD3^+^, DAPI^+^, PanCK-) were manually counted in stroma and tumor areas. Final counts were normalized to total area of stroma or tumor annotation.

### IVIS imaging

#### In vivo

Mice were injected with 100 uL of 15 mg/mL D-luciferin potassium salt in DPBS (no Ca, no Mg) (GoldBio, #LUCK-100), then anesthetized with isoflurane. 10 minutes later, a luminescent image and photo were captured. Longitudinal data was analyzed in one batch by normalizing tumor ROI luminescence total counts to average background ROI luminescence total counts.

#### Ex vivo

Organs for analysis were placed in PBS on ice immediately following dissection. When all organs were collected, PBS was replaced with 1 mL of 15 mg/mL D-Luciferin in DPBS (no Ca, no Mg) (GoldBio, #LUCK-100). Organs were incubated for 5 minutes before removing from solution, patting dry, and arranging on the IVIS imaging tray. Luminescent image and photo were captured. Total luminescence counts were normalized to background luminescence counts.

### Lipid nanoparticle (LNP) preparation

LNPs were prepared using microfluidic mixing of ethanol (lipid) and aqueous (mRNA) phases at a 3:1 volumetric ratio, with a total flow rate of 12 mL per minute using the Ignite system (Precision Nanosystems, BC, Canada). The lipid phase contained ionizable lipid SM102, cholesterol, DSPC, and DMG-PEG2000 at 50/10/38.5/1.5 molar ratios, respectively, diluted in 100% ethanol. mRNA was diluted in sterile 50 mM citrate buffer to achieve an N/P ratio of 5.67. For empty LNPs, an equivalent amount of citrate buffer was used. After aqueous and lipid solutions were mixed, the LNPs were dialyzed for 4 hours at room temperature, followed by an overnight dialysis against sterile PBS at 4C in 10 kDa Slide-a-Lyzer G3 cassettes (ThermoFisher, MA, USA). Then, LNPs were concentrated by centrifugation using 100k MWCO Amicon ultracentrifugal filters (Millipore Sigma, Burlington, MA) at 3000g and 4C. LNPs were stored at 4C before administration.

### LNP Characterization

Hydrodynamic size and polydispersity of the LNPs were determined with DLS using Stunner (Unchained Labs, Pleasanton, CA). Encapsulation efficiency and concentration of mRNA were determined using a modified Quant-iT RiboGreen RNA assay (ThermoFisher, MA, USA) and a multimode microplate reader. Cholesterol content was determined with the Invitrogen™ Amplex™ Red Cholesterol Assay Kit (ThermoFisher, MA, USA).

### Plasma analytes

Plasma was collected, snap frozen in liquid nitrogen, and stored at -80°C. Plasma concentrations of IL-6 (Biolegend) were measured using ELISA, and read on a plate reader (BioTek).

### Quantitative real-time polymerase chain reaction (qPCR)

We isolated RNA from cell pellets or tissue samples using the E.Z.N.A. Total RNA Kit I (Omega BioTek) and we prepared cDNA using high-capacity cDNA reverse transcription kit (Applied Biosystems). qPCR was run on the ABI 7300 (Applied Biosystems), using TaqMan Fast Advanced PCR master mix (Applied Biosystems) or SYBR Green master mix (Applied Biosystems). Relative expression was calculated using the ΔΔC_t_ method. To confirm the presence/absence of *Il6 transgene* in tumor-implanted mice, we performed 30 cycles of qPCR on the ABI7300, followed by running the PCR product on a 3% ethidium bromide gel to determine the presence of a band at the expected size of 131 bp. Primers/probes are listed in **Table S2**.

### Statistical Analysis

Specific statistical tests and sample size for each study is indicated in the figure legends. Error bars in figures show SEM. Statistical analyses were performed using GraphPad Prism (version 9; GraphPad Software Inc) or JMP Pro (version 16; SAS Institute Inc), and graphs were built using GraphPad Prism (GraphPad Software Inc) statistical analysis software. P values are 2 sided with values less than 0.05 regarded as statistically significant.

### Data Availability

All authors had access to the study data and had reviewed and approved the final manuscript. All data will be available on the public repository, Mendeley Data, upon publication. Further information and resources, including plasmid sequences, engineered KPC cells, and raw data will be shared upon reasonable request to Aaron J. Grossberg.

## ACKNOWLEDGEMENTS

We thank all members of the Aaron Grossberg, Robert Eil, and Katelyn Byrne, and Gaurav Sahay labs for their helpful discussion and suggestions. We also acknowledge the expert technical assistance by staff in the Advanced Multiscale Microscopy Shared Resource and Histopathology Shared Resource. Author contributions are: Conceptualization, PCAW, AQB, RE, AJG. Methodology, PCAW, AQB, NTVM, AJ, YE, GS, KTB, RE, AJG. Validation, PCAW, AQB, HM, XZ, JD, MM, PRL, PD. Formal Analysis, PCAW, AQB, HM, XZ, JD, PD, GDS, AJG. Investigation, PCAW, AQB, HM, XZ, JD, MM, PRL, PD, GDS. Writing—Original Draft, PCAW. Writing – Review and Editing, PCAW, AQB, KTB, RE, AJG. Visualization, PCAW, AQB, PD. Supervision, RE, AJG. Project Administration, RE, AJG. Funding Acquisition, PCAW, RE, AJG. All authors approved this manuscript.

## Data Transparency

All data will be available on the public repository, Mendeley Data, upon publication. Further information and resources, including plasmid sequences, engineered KPC cells, and raw data will be shared upon reasonable request to Aaron J. Grossberg.

## Disclosures

GS is a co-founder of EnterX Bio and has an advisory role to Rare Air Inc., Serina Tx, and Mana Bio. KTB receives consultation feed from Guidepoint Global and royalties from the University of Pennsylvania for licensed research cell lines. The other authors declare no conflicts. RE is a paid consultant and conducts ongoing research for Lyell Immunopharma. AJG is a paid consultant for Endevica Bio, Inc.

### Abbreviations

BSR: Blasticidin Resistance
CAR: Chimeric Antigen Receptor
CD: Cell Differentiation
ELISA: Enzyme-Linked Immunosorbent Assay
Fbxo32: F-Box Protein 32
Foxp3: Forkhead Box Protein P3
H&E: Hematoxylin and Eosin
IL-6: Interleukin 6
KPC: Kras^G12D/+^, Tp53^R172H/+,^ Pdx1-Cre
NK: Natural Killer
OT-PDAC: Orthotopic Pancreatic Ductal Adenocarcinoma
PDAC: Pancreatic Ductal Adenocarcinoma
Qpcr: Quantitative Polymerase Chain Reaction
Trim63: Tripartite Motif Containing 63

---

**Figure S1:**
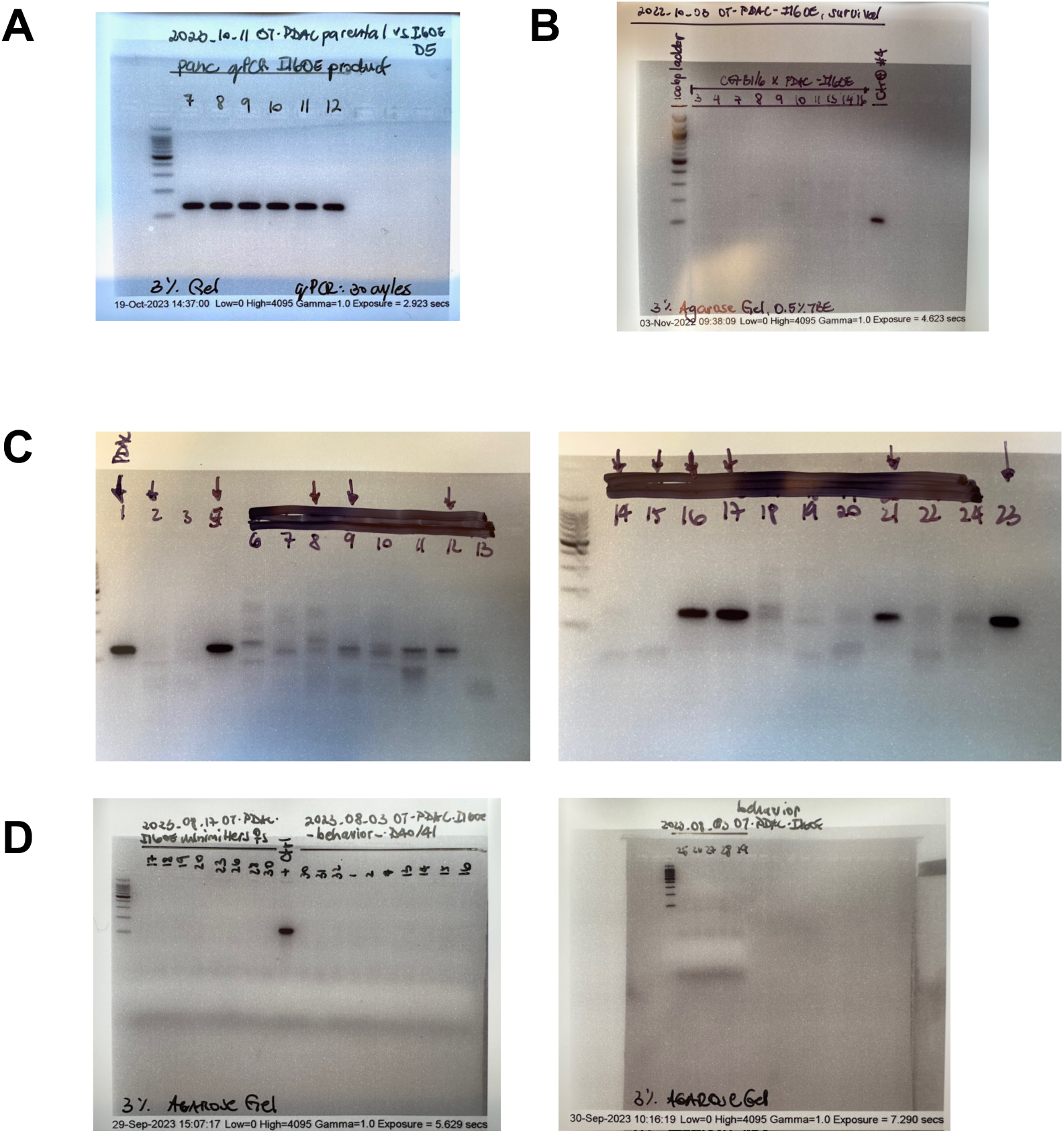
OT-KxPxCx^IL6^ clearance detected by *Il6-transgene* qPCR product. qPCR products from pancreas/tumor tissue, run for 30 PCR cycles, and then run out on a 3% agarose gel. Expected product is 131 bp. Ladder is Promega 100 bp ladder (G210A). (A) 5 days post implantation. (B) 17 days post implantation (samples 7, 9, 11, 16) and 24 days post implantation (samples 3, 4, 8, 10, 14). (C)10 days post implantation. (D) 40 days post implantation.

**Figure S2:**
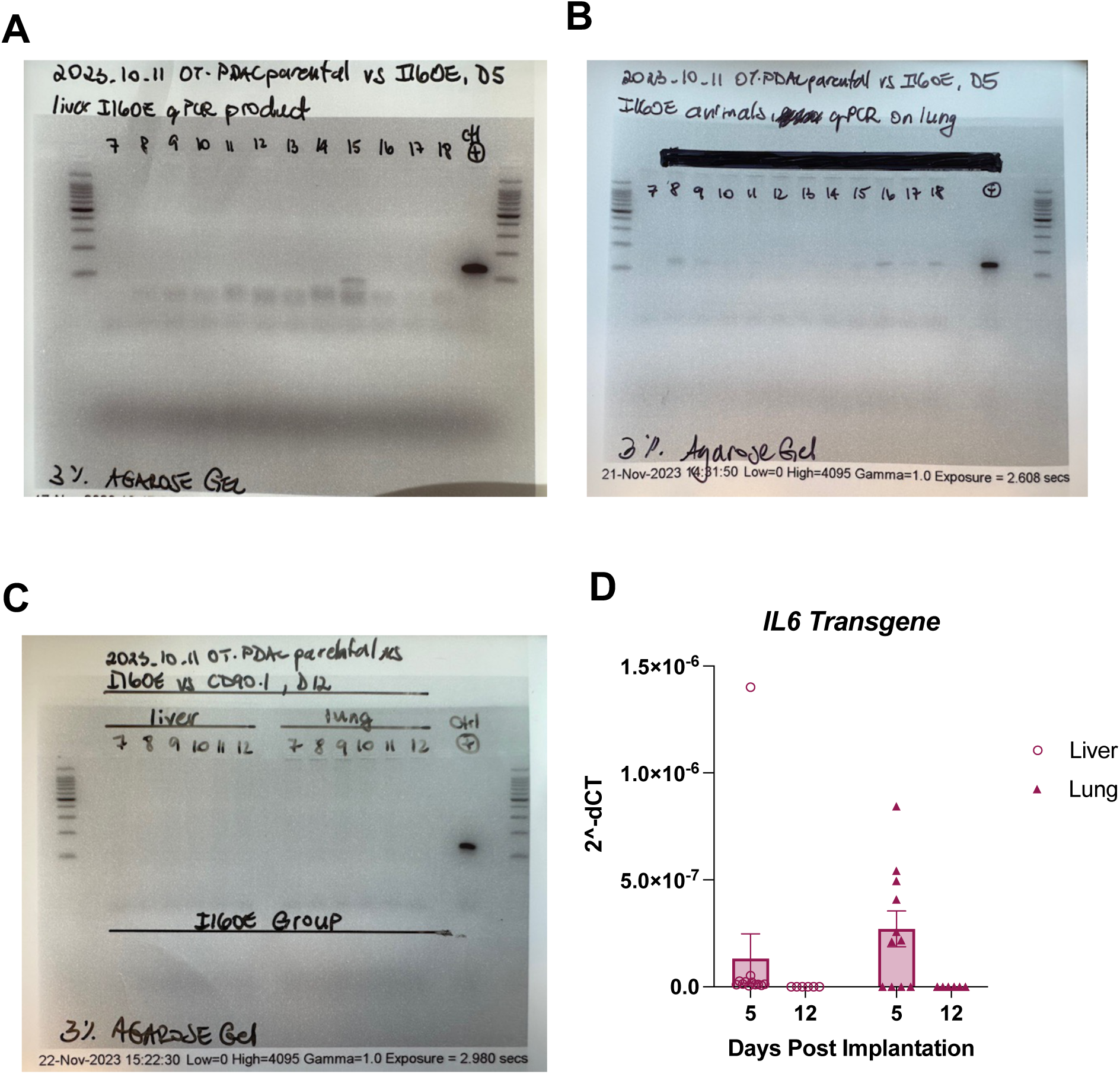
KxPxCx^IL6^ metastatic cell presence detected by *Il6-transgene* qPCR product. qPCR products, run for 30 PCR cycles, and then run out on a 3% agarose gel. Expected product is 131 bp. Ladder is Promega 100 bp ladder (G210A). (A) Whole liver at 5 days post implantation. (B) Whole lung at 5 days post implantation. (C) Whole lung and liver at 12 days post implantation. (D) Quantification.

**Figure S3:**
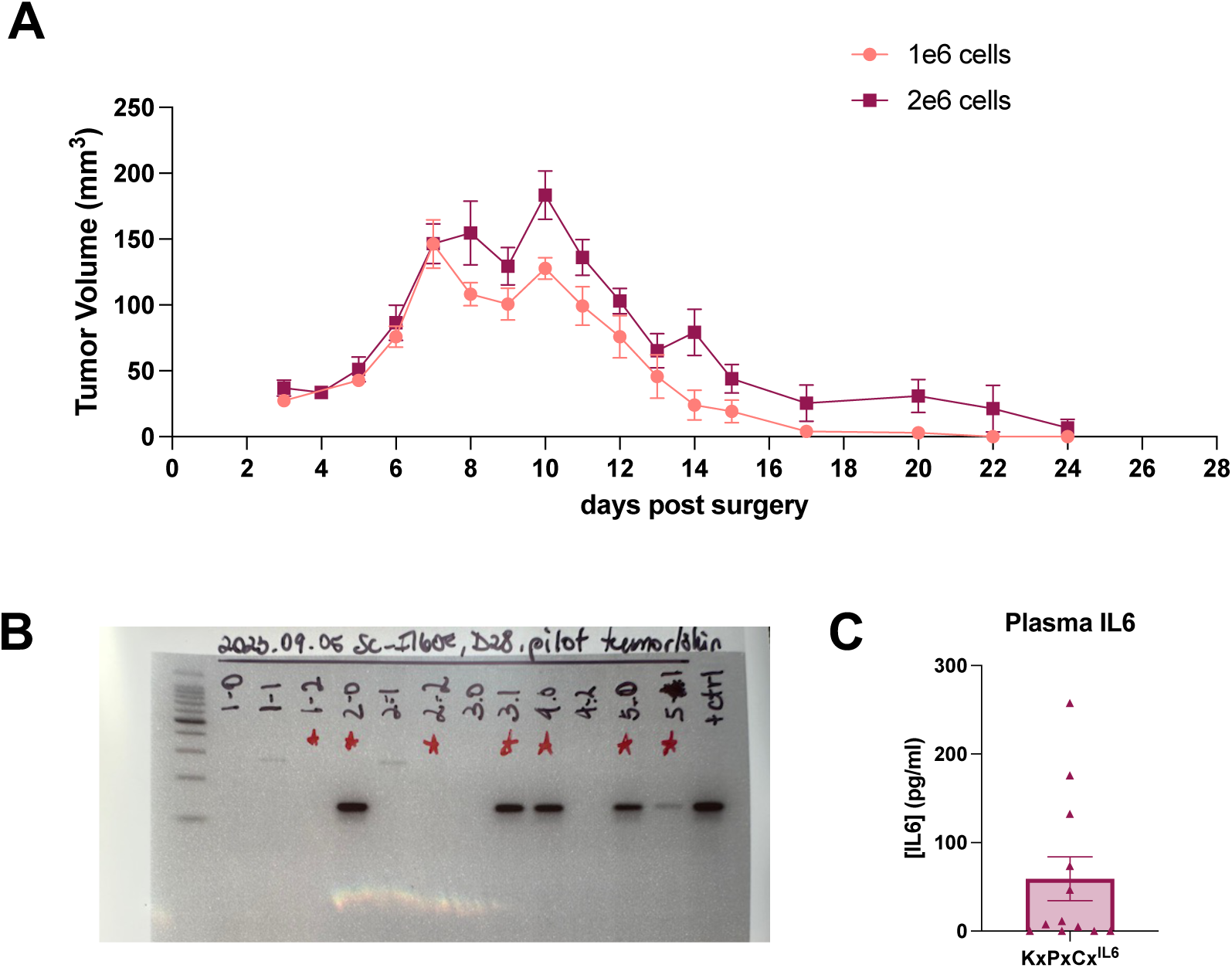
KxPxCx^IL6^ cells induce anti-tumor immune response when implanted subcutaneously. (A) Tumor volume measured by calipers daily. (B) Injection site/tumor tissue collected at 24 days post implantation qPCR products, run for 30 PCR cycles, and then run out on a 3% agarose gel. Expected product is 131 bp. Ladder is Promega 100 bp ladder (G210A). (C) Plasma IL-6 measured by ELISA at 24 days post implantation.

**Figure S4:**
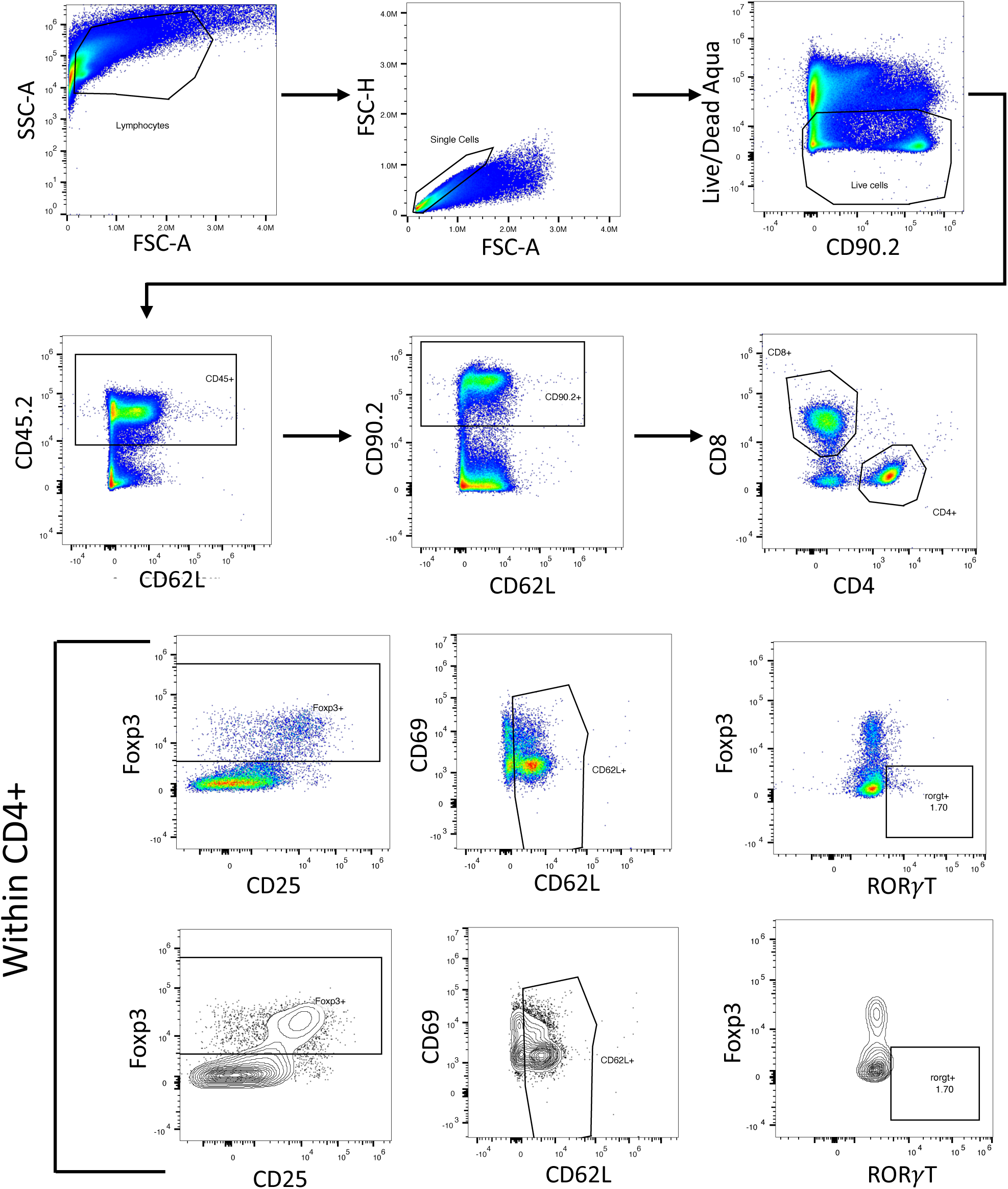
Gating strategy for flow cytometry presented in Figure 4B-G, and Figure S6 B-E, G (representative sample).

**Figure S5:**
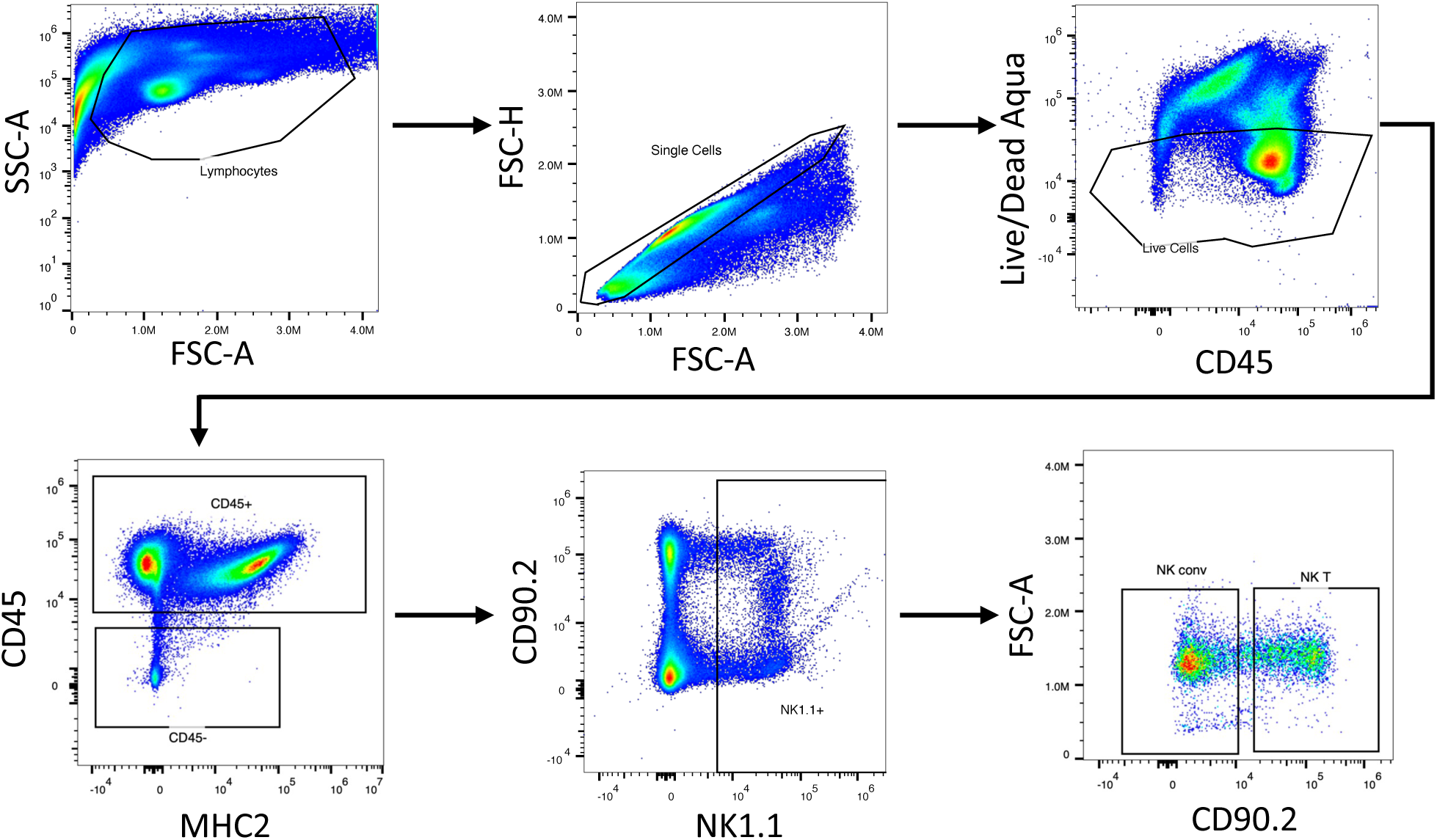
Gating strategies for flow cytometry presented in Figure 4H-I, and Figure S6A, F (representative sample).

**Figure S6:**
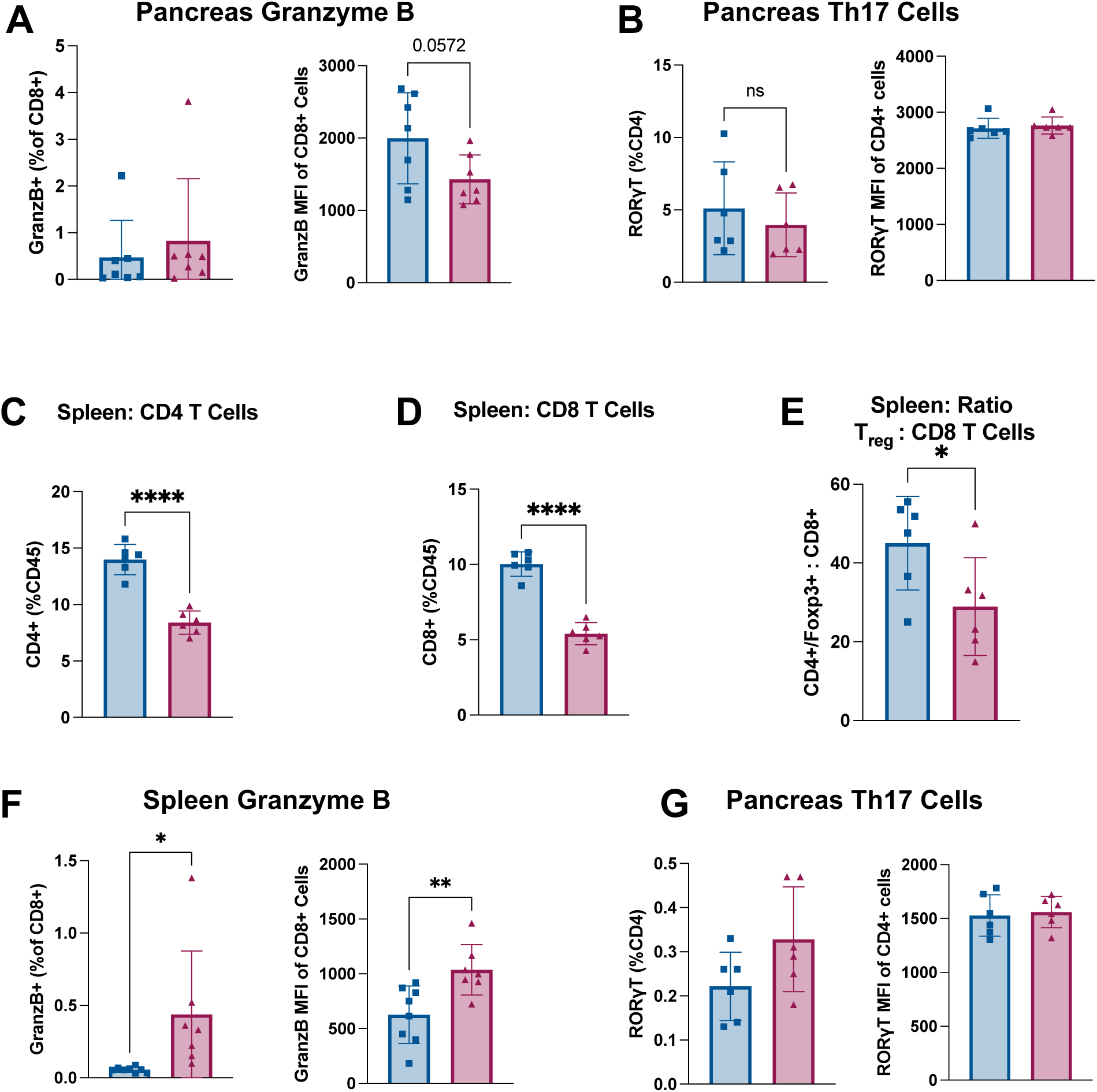
Supporting flow cytometry data for Th17 and Granzyme B+ T cell phenotypes. Intra-tumoral immune cell populations from pancreas at 5 days: (A) Granzyme B+ cells as a percent of CD8+ cells (left) and CD8+ Granzyme B MFI (right). (B) Th17 Cells (ROR!T+) as a percent of CD4+ cells (left) and CD4+ ROR!T MFI (right). Immune cell populations from spleen at 5 days: (C) CD4^+^ T cells, (D) CD8^+^ T cells, (E) Ratio of Foxp3^+^ to CD8^+^ cells, (F) Granzyme B+ cells as a percent of CD8+ cells (left) and CD8+ Granzyme B MFI (right). (G) Th17 Cells (ROR!T+) as a percent of CD4+ cells (left) and CD4+ ROR!T MFI (right). (B-E, G) N = 6 male mice per group. (A, F) N = 4 female, 3 male mice per group. Error bars represent SEM. 2-group analysis tested with unpaired t-test. **** p<0.0001, ***p<0.001, **p<0.01, *p<0.05.

**Figure S7:**
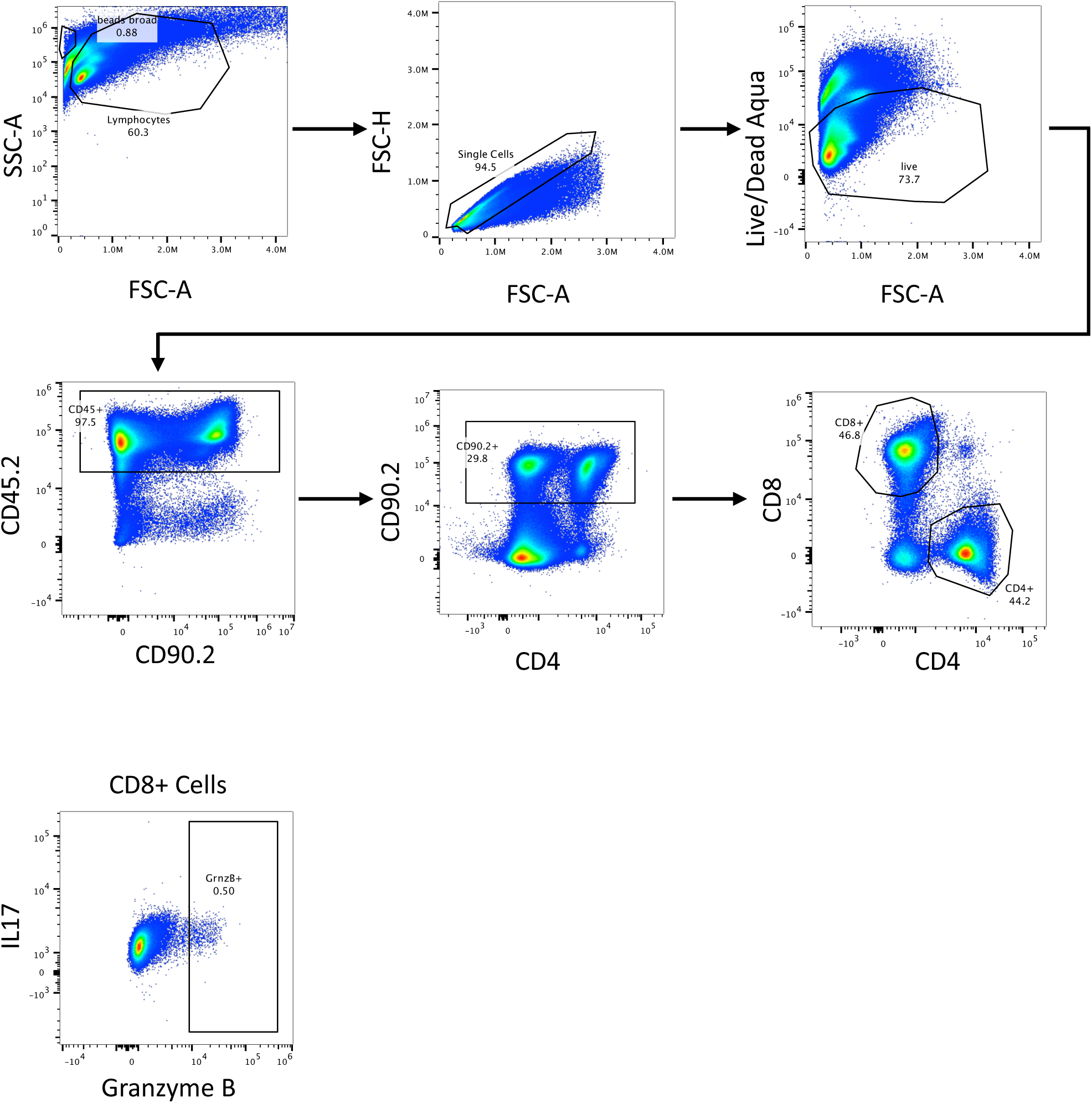
Granzyme B gating strategy for flow cytometry presented in Figure S6 (representative samples).

**Figure S8:**
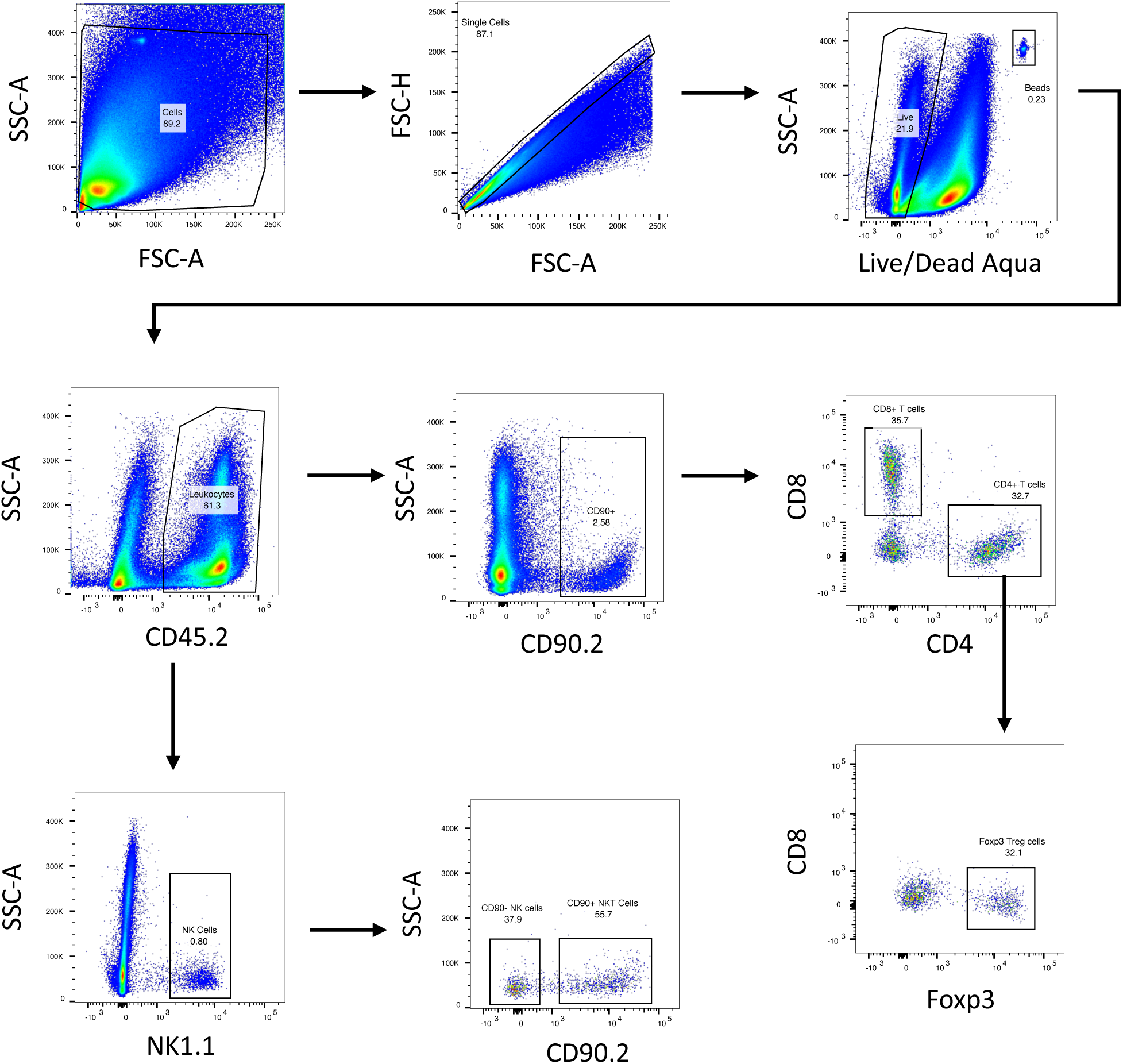
Gating strategies for flow cytometry presented in Figure 7 (representative sample).

**Figure S9:**
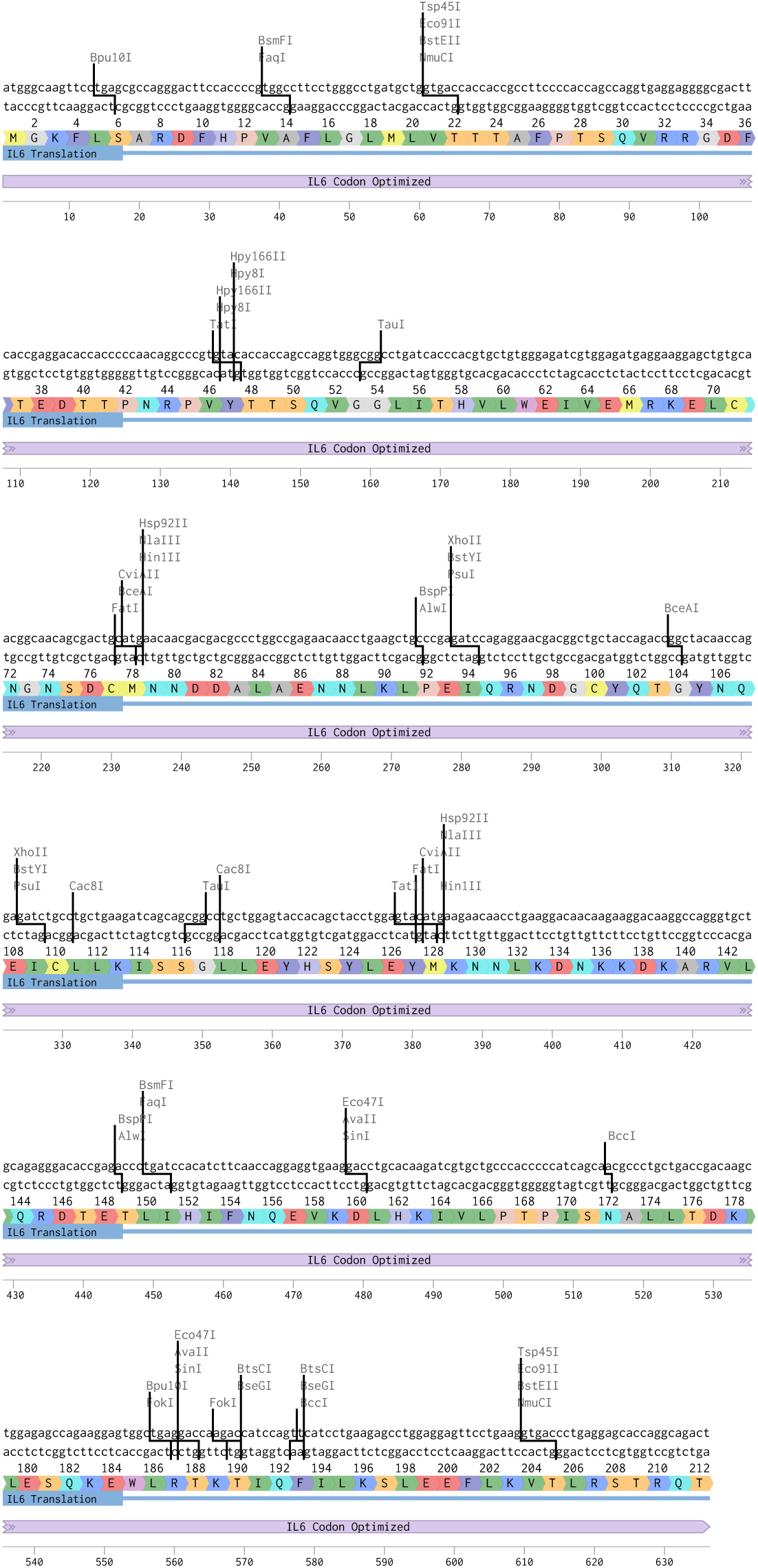
Linear sequence map with amino acid annotation for codon-optimized IL6 overexpression cassette transduced into PDAC cells.

**Table S1:**
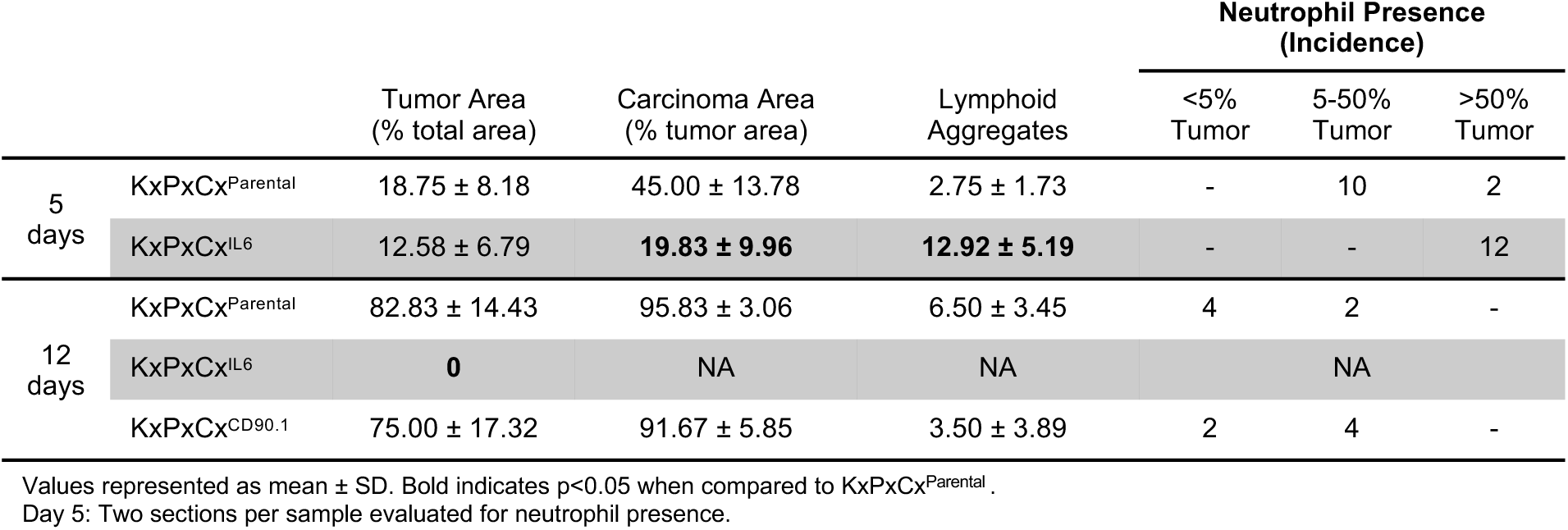
Histological assessment of transgenic tumors. .

**Table S2:**
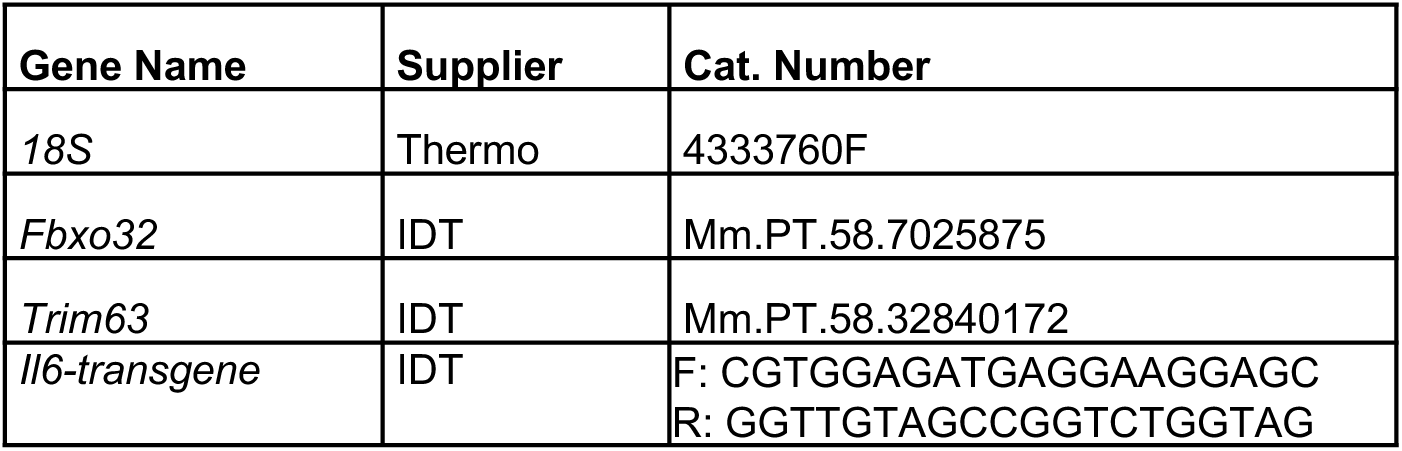
qPCR primer and probe information.

**Table S3:**
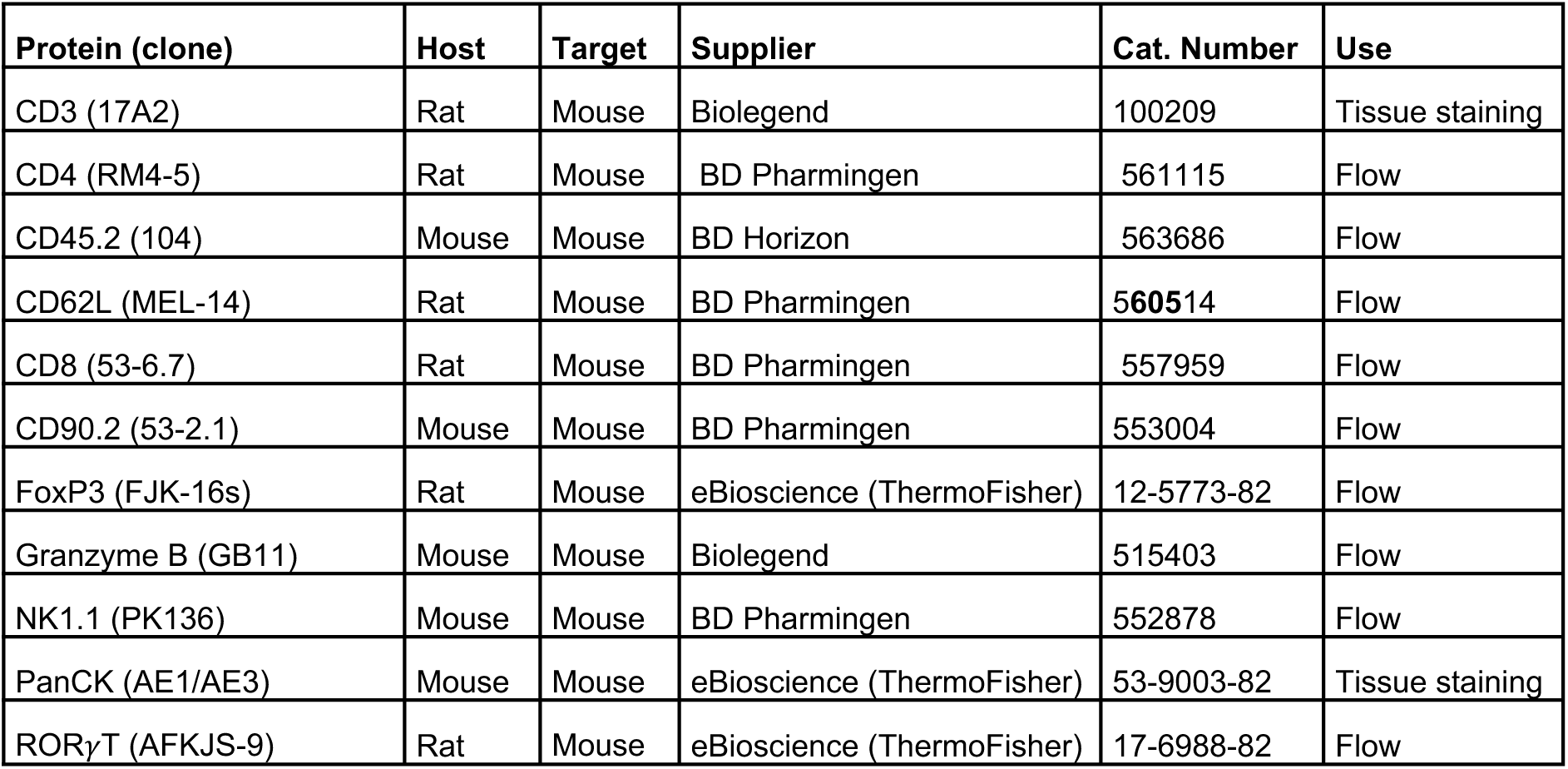
Flow cytometry antibody information.

