## supplemental information for "Interleukin 6 drives durable T cell-mediated immunity to pancreatic cancer"

Table S1: Histological assessment of transgenic tumors.

| | | Tumor Area (% total area) | Carcinoma Area (% tumor area) | Lymphoid Aggregates | Neutrophil Presence (Incidence) | | |
|---|---|---|---|---|---|---|---|
| | | | | | <5% Tumor | 5-50% Tumor | >50% Tumor |
| 5 days | KxPxCx$^{Parental}$ | 18.75 ± 8.18 | 45.00 ± 13.78 | 2.75 ± 1.73 | - | 10 | 2 |
| | KxPxCx$^{IL6}$ | 12.58 ± 6.79 | **19.83 ± 9.96** | **12.92 ± 5.19** | - | - | 12 |
| 12 days | KxPxCx$^{Parental}$ | 82.83 ± 14.43 | 95.83 ± 3.06 | 6.50 ± 3.45 | 4 | 2 | - |
| | KxPxCx$^{IL6}$ | **0** | NA | NA | NA | | |
| | KxPxCx$^{CD90.1}$ | 75.00 ± 17.32 | 91.67 ± 5.85 | 3.50 ± 3.89 | 2 | 4 | - |

Values represented as mean ± SD. Bold indicates $p<0.05$ when compared to KxPxCx$^{Parental}$ .
Day 5: Two sections per sample evaluated for neutrophil presence.

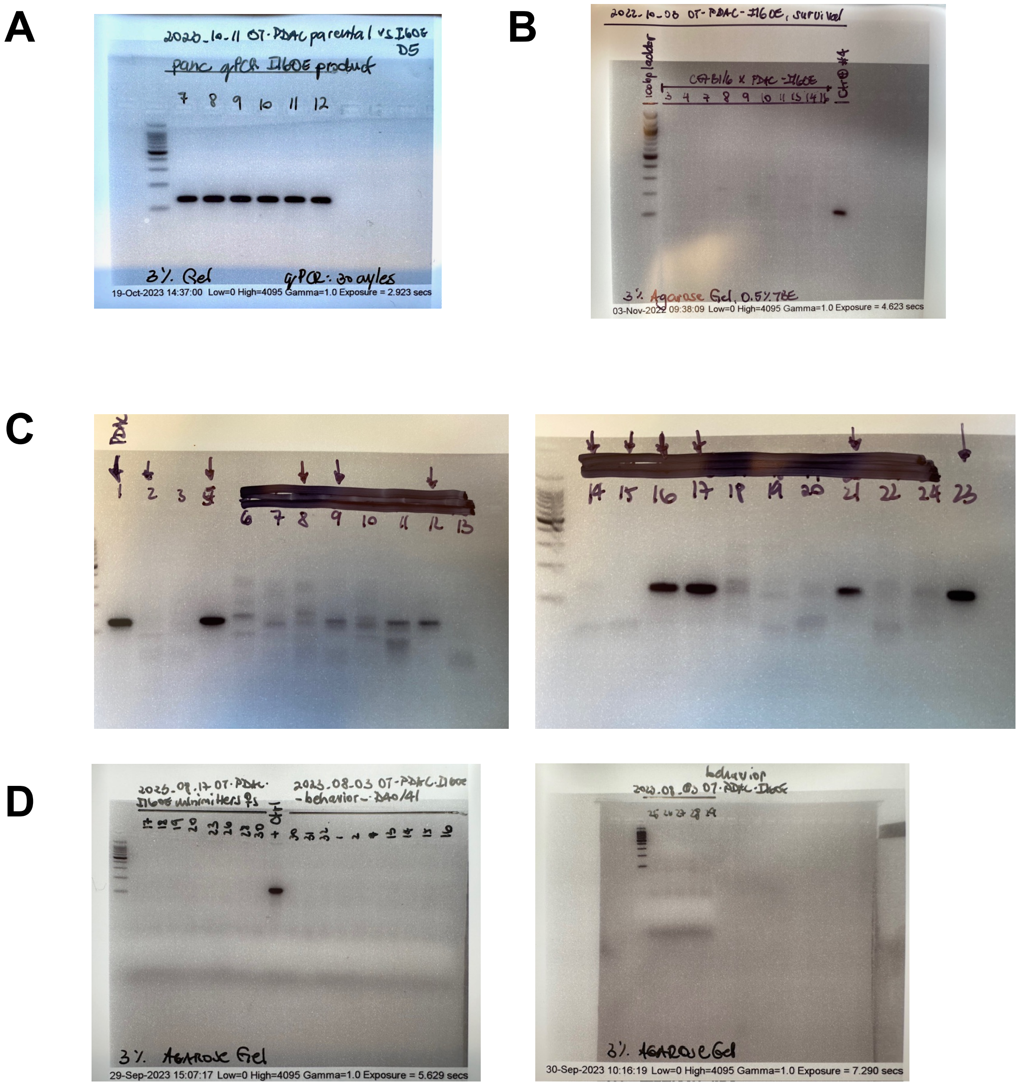


**Figure S1**: OT-KxPxCx$^{IL6}$ clearance detected by *Il6-transgene* qPCR product. qPCR products from pancreas/tumor tissue, run for 30 PCR cycles, and then run out on a 3% agarose gel. Expected product is 131 bp. Ladder is Promega 100 bp ladder (G210A). (A) 5 days post implantation. (B) 17 days post implantation (samples 7, 9, 11, 16) and 24 days post implantation (samples 3, 4, 8, 10, 14). (C)10 days post implantation. (D) 40 days post implantation.

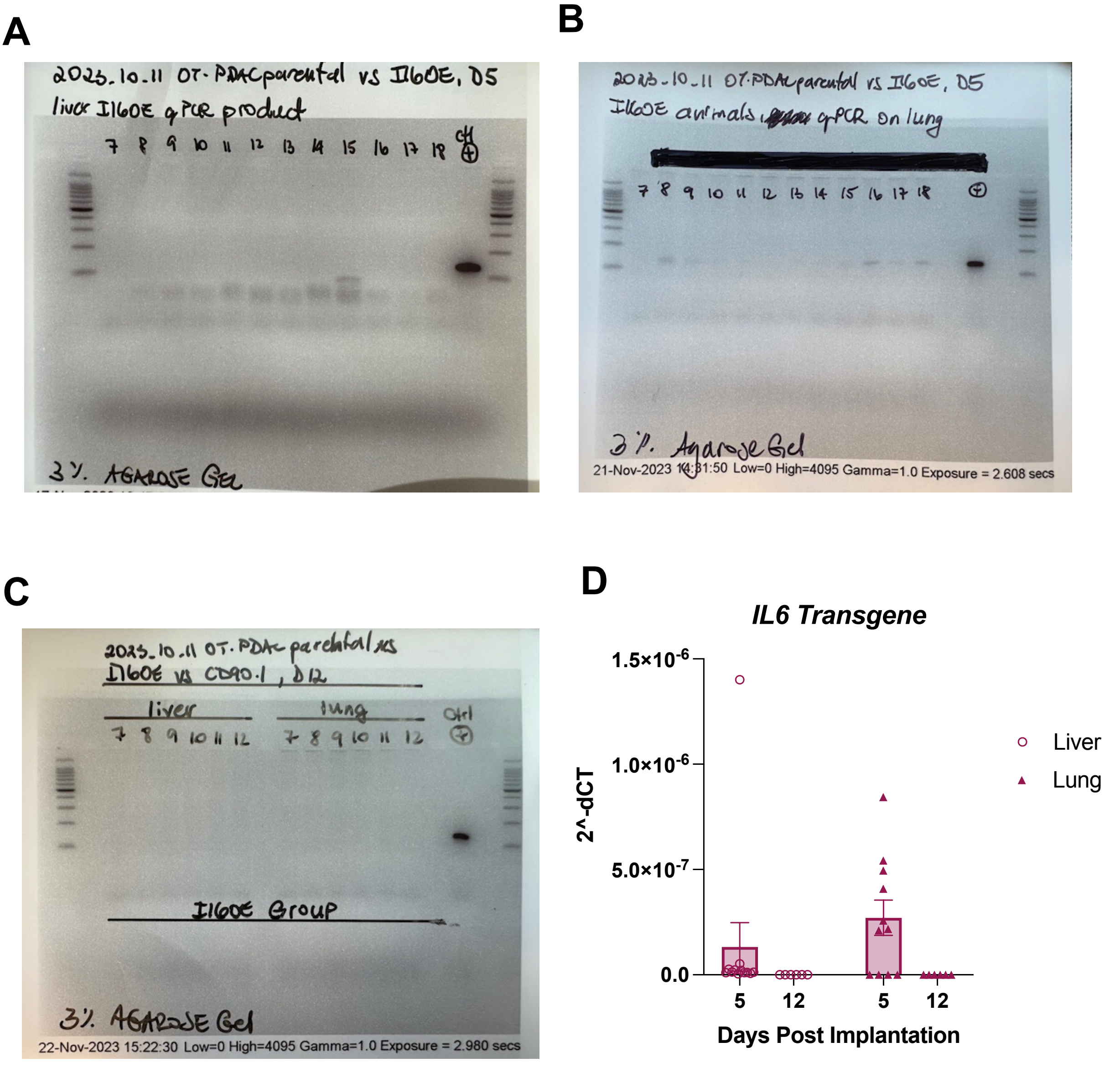


**Figure S2**: KxPxCx$^{IL6}$ metastatic cell presence detected by *Il6-transgene* qPCR product. qPCR products, run for 30 PCR cycles, and then run out on a 3% agarose gel. Expected product is 131 bp. Ladder is Promega 100 bp ladder (G210A). (A) Whole liver at 5 days post implantation. (B) Whole lung at 5 days post implantation. (C) Whole lung and liver at 12 days post implantation. (D) Quantification.

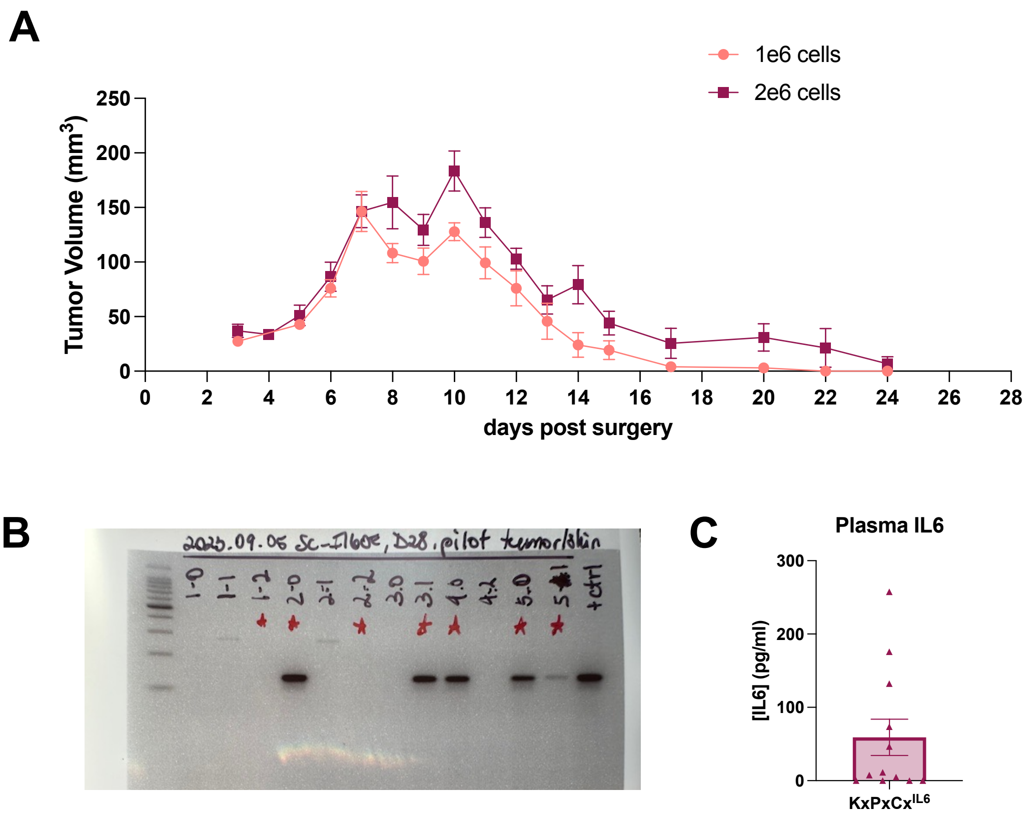


**Figure S3**: KxPxCx$^{IL6}$ cells induce anti-tumor immune response when implanted subcutaneously. (A) Tumor volume measured by calipers daily. (B) Injection site/tumor tissue collected at 24 days post implantation qPCR products, run for 30 PCR cycles, and then run out on a 3% agarose gel. Expected product is 131 bp. Ladder is Promega 100 bp ladder (G210A). (C) Plasma IL-6 measured by ELISA at 24 days post implantation.

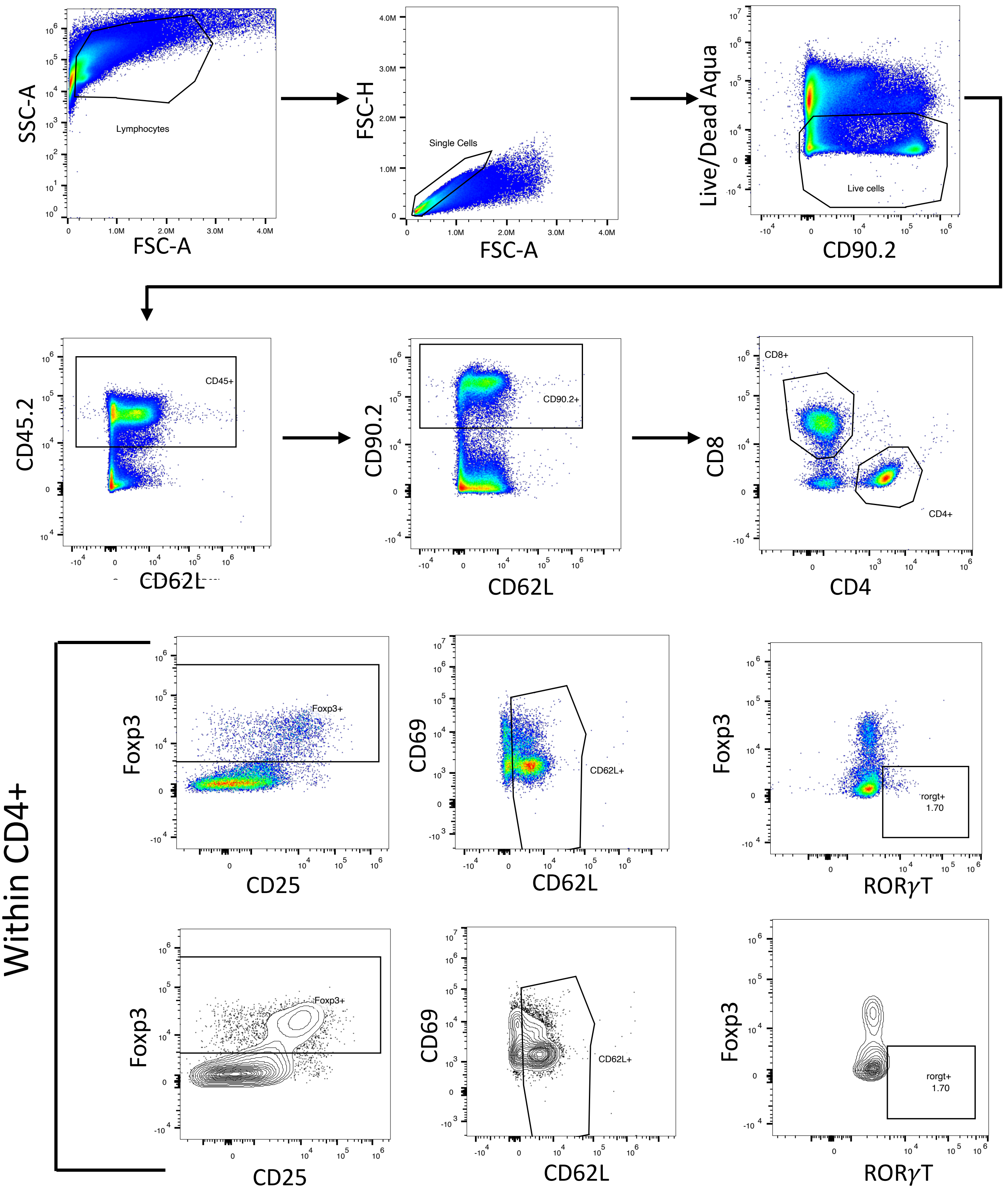


**Figure S4**: Gating strategy for flow cytometry presented in Figure 4B-G, and Figure S6 B-E, G (representative sample).

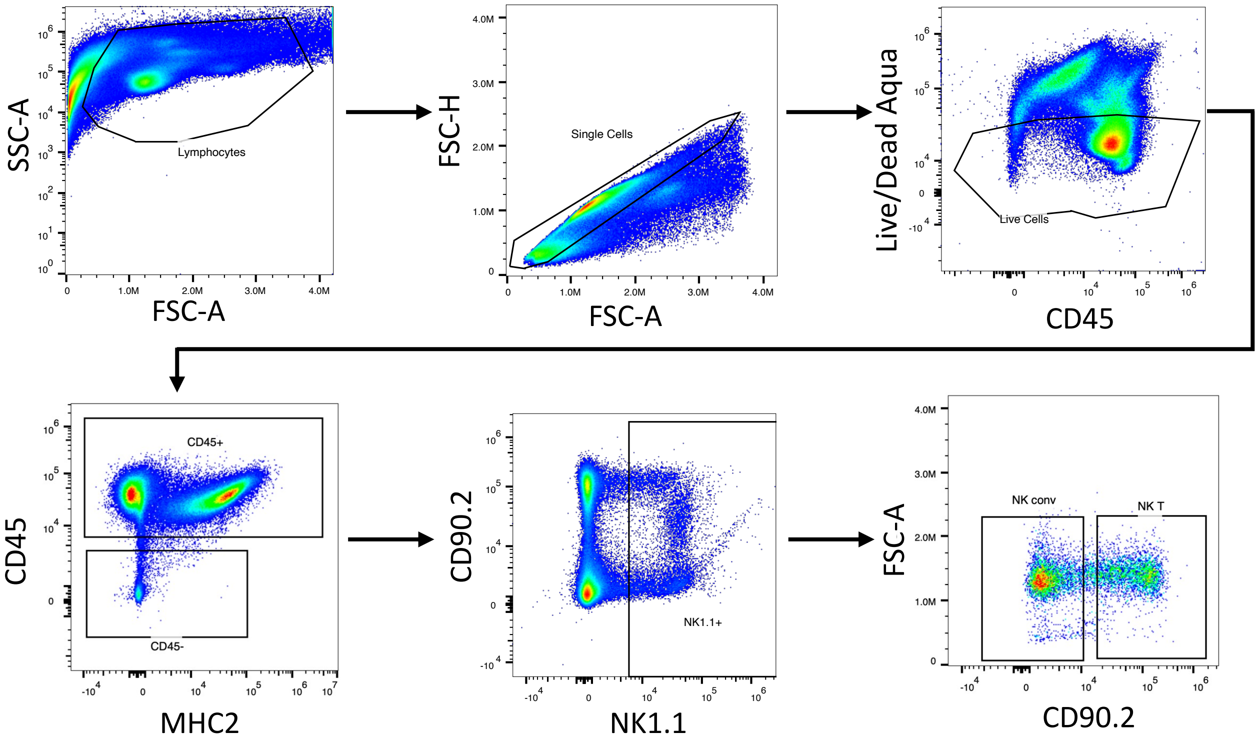


**Figure S5**: Gating strategies for flow cytometry presented in Figure 4H-I, and Figure S6A, F (representative sample).

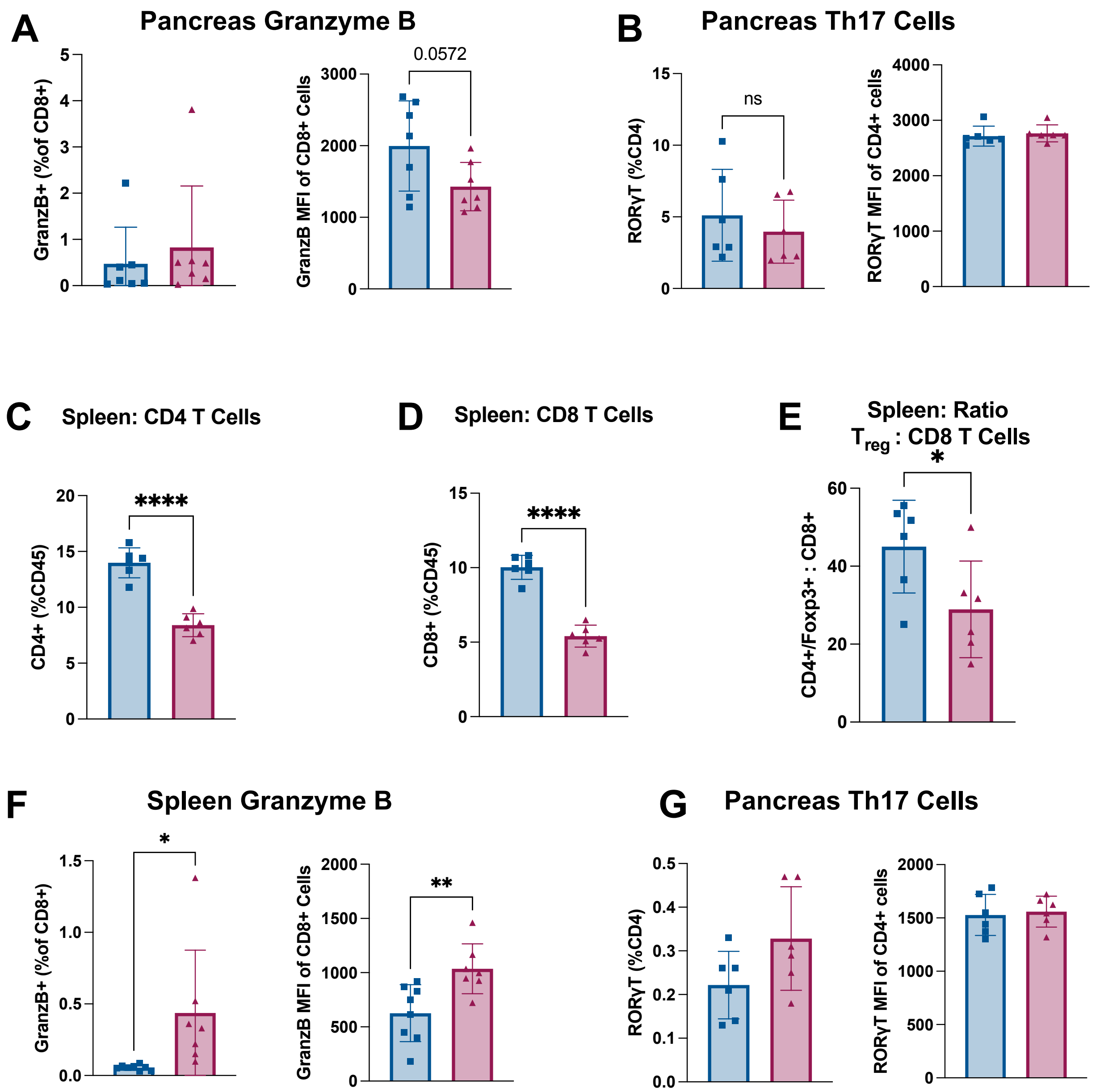


**Figure S6:** Supporting flow cytometry data for Th17 and Granzyme B+ T cell phenotypes. Intra-tumoral immune cell populations from pancreas at 5 days: (A) Granzyme B+ cells as a percent of CD8+ cells (left) and CD8+ Granzyme B MFI (right). (B) Th17 Cells (ROR$\gamma$T+) as a percent of CD4+ cells (left) and CD4+ ROR$\gamma$T MFI (right). Immune cell populations from spleen at 5 days: (C) $CD4^+$ T cells, (D) $CD8^+$ T cells, (E) Ratio of $Foxp3^+$ to $CD8^+$ cells, (F) Granzyme B+ cells as a percent of CD8+ cells (left) and CD8+ Granzyme B MFI (right). (G) Th17 Cells (ROR$\gamma$T+) as a percent of CD4+ cells (left) and CD4+ ROR$\gamma$T MFI (right). (B-E, G) N = 6 male mice per group. (A, F) N = 4 female, 3 male mice per group. Error bars represent SEM. 2-group analysis tested with unpaired t-test. **** $p<0.0001$, ***$p<0.001$, **$p<0.01$, *$p<0.05$.

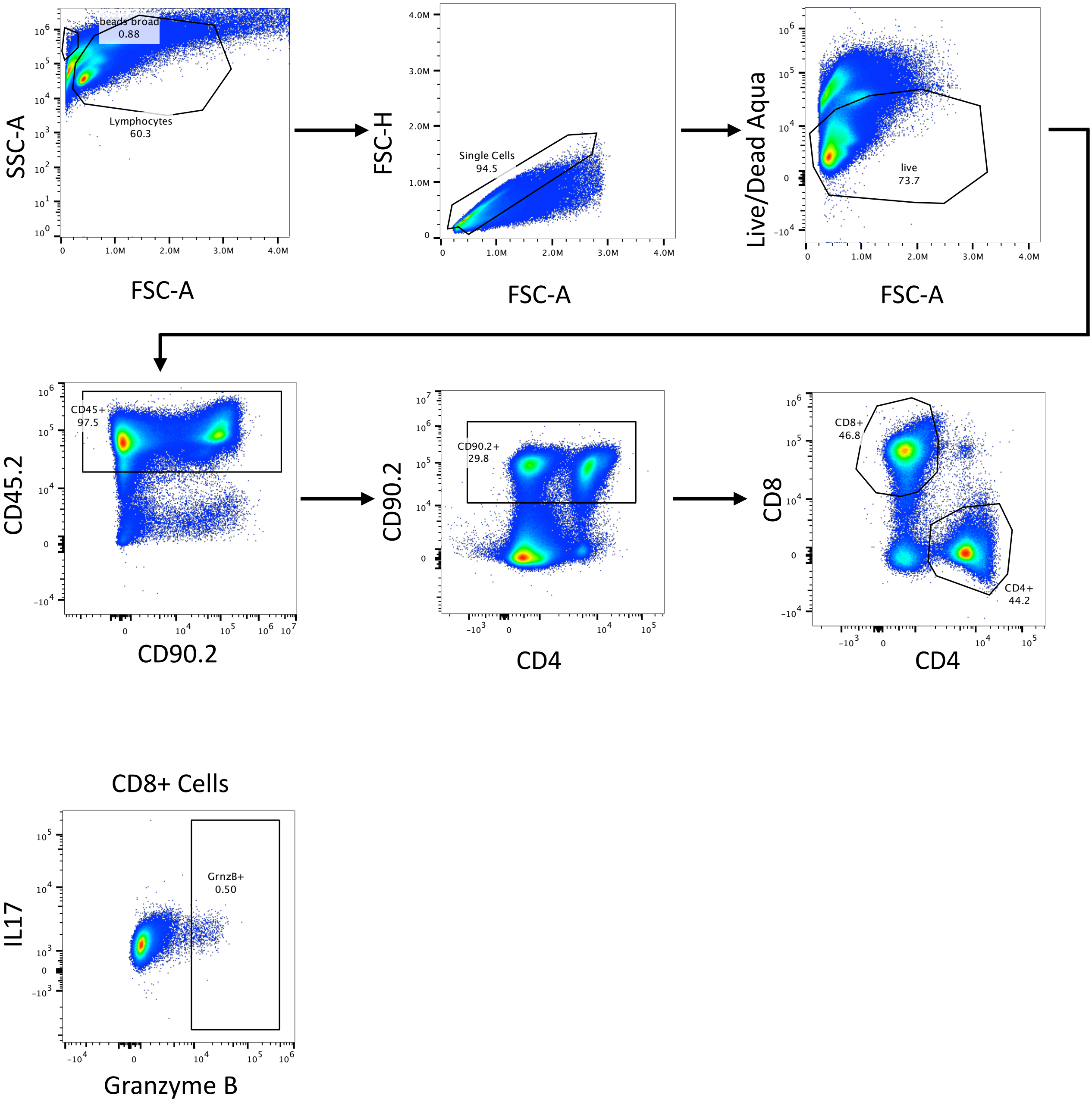


**Figure S7**: Granzyme B gating strategy for flow cytometry presented in Figure S6 (representative samples).

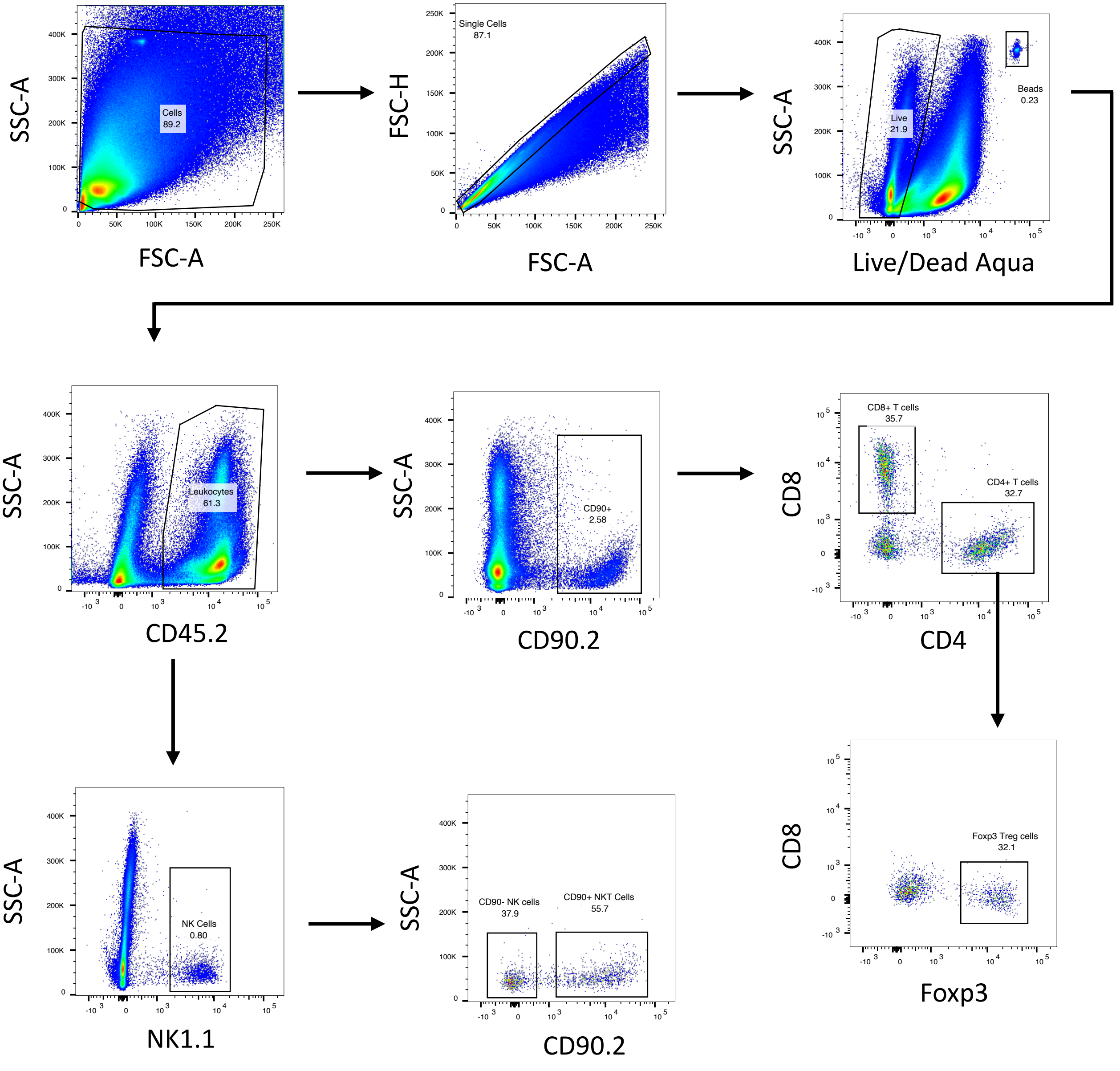


**Figure S8**: Gating strategies for flow cytometry presented in Figure 7 (representative sample).

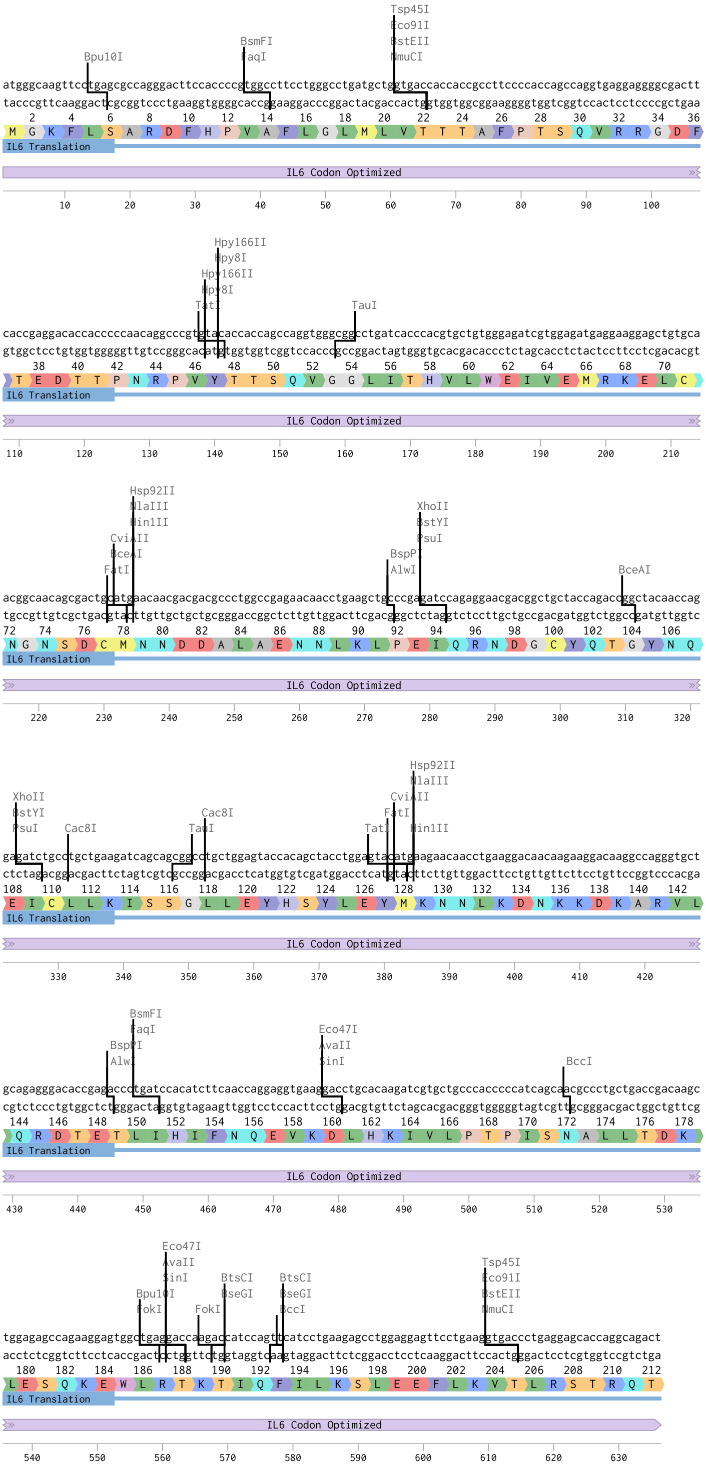


**Figure S9**: Linear sequence map with amino acid annotation for codon-optimized IL6 overexpression cassette transduced into PDAC cells.

Table S2: qPCR primer and probe information

| Gene Name | Supplier | Cat. Number |
|---|---|---|
| *18S* | Thermo | 4333760F |
| *Fbxo32* | IDT | Mm.PT.58.7025875 |
| *Trim63* | IDT | Mm.PT.58.32840172 |
| *Il6-transgene* | IDT | F: CGTGGAGATGAGGAAGGAGC<br>R: GGTTGTAGCCGGTCTGGTAG |

Table S3: Flow cytometry antibody information

| **Protein (clone)** | **Host** | **Target** | **Supplier** | **Cat. Number** | **Use** |
|---|---|---|---|---|---|
| CD3 (17A2) | Rat | Mouse | Biolegend | 100209 | Tissue staining |
| CD4 (RM4-5) | Rat | Mouse | BD Pharmingen | 561115 | Flow |
| CD45.2 (104) | Mouse | Mouse | BD Horizon | 563686 | Flow |
| CD62L (MEL-14) | Rat | Mouse | BD Pharmingen | 5**605**14 | Flow |
| CD8 (53-6.7) | Rat | Mouse | BD Pharmingen | 557959 | Flow |
| CD90.2 (53-2.1) | Mouse | Mouse | BD Pharmingen | 553004 | Flow |
| FoxP3 (FJK-16s) | Rat | Mouse | eBioscience (ThermoFisher) | 12-5773-82 | Flow |
| Granzyme B (GB11) | Mouse | Mouse | Biolegend | 515403 | Flow |
| NK1.1 (PK136) | Mouse | Mouse | BD Pharmingen | 552878 | Flow |
| PanCK (AE1/AE3) | Mouse | Mouse | eBioscience (ThermoFisher) | 53-9003-82 | Tissue staining |
| ROR$\gamma$T (AFKJS-9) | Rat | Mouse | eBioscience (ThermoFisher) | 17-6988-82 | Flow |
